# Fusion-driven oncogenic programs shape the immune landscape in translocation renal cell carcinoma

**DOI:** 10.1101/2025.08.28.672849

**Authors:** Prathyusha Konda, Cary N Weiss, Yantong Cui, Sayed Matar, Jinyu Wang, Yasmin L. Nabil, Riva Deodhar, Jiao Li, Jack Horst, Sabrina Y. Camp, Aseman B. Sheshdeh, Jonathan L. Hecht, David J. Einstein, Yashika Rustagi, Anwesha Nag, Aaron R. Thorner, Cheng-Zhong Zhang, Eliezer M. Van Allen, Sabina Signoretti, Toni K. Choueiri, Srinivas R. Viswanathan

**Affiliations:** Department of Medical Oncology, Dana-Farber Cancer Institute, Boston, MA 02215, USA; Department of Medicine, Harvard Medical School, Boston, MA 02215, USA; Department of Pediatric Oncology, Dana-Farber Cancer Institute, Boston, MA 02215, USA; Department of Pathology, Brigham and Women’s Hospital, Boston, MA 02215, USA; Department of Biomedical Informatics, Dana-Farber Cancer Institute, Boston, MA 02215, USA; Department of Pathology, Beth Israel Deaconess Medical Center, Boston, MA 02215, USA; Division of Medical Oncology, Beth Israel Deaconess Medical Center, Boston, MA 02215, USA; Broad Institute of MIT and Harvard, Cambridge, MA 02142, USA; Department of Medicine, Brigham and Women’s Hospital; Boston, MA 02215, USA

## Abstract

Renal cell carcinomas comprise multiple molecularly distinct cancers but most are treated empirically with therapies designed for clear cell RCC (ccRCC), the most common subtype, due to incomplete understanding of subtype-specific biology. We analyzed single-cell transcriptomes and chromatin accessibility profiles from translocation renal cell carcinoma (tRCC), an aggressive RCC defined by oncogenic *TFE3* gene fusions. Unexpectedly, despite arising from a proximal tubule cell of origin similar to ccRCC, tRCCs display markedly distinct oncogenic programs and an immunosuppressive tumor microenvironment. tRCCs exhibit six conserved tumor meta-programs, including epithelial-mesenchymal transition and proximal tubule identity programs whose balance is regulated by TFE3 fusion activity. The fusion-driven EMT program drives a suppressive TME marked by progenitor-exhausted CD8+ T cells, anti-inflammatory SPP1+ macrophages, and matrix-associated fibroblasts (mCAFs). Our findings highlight unique TFE3 fusion-driven biology in tRCC, explaining its reduced immunotherapy responsiveness relative to ccRCC, and suggesting strategies for targeting fusion-driven oncogenic programs and TME reprogramming.

## INTRODUCTION

Kidney cancer comprises dozens of different subtypes, each with unique molecular and clinical characteristics. To date, most discovery biology research in kidney cancer has focused on clear-cell renal cell carcinoma (ccRCC), which constitutes approximately 75% of renal cell carcinomas in adults^1,2^. ccRCC is characterized by a pathognomonic inactivation of the Von Hippel-Lindau (*VHL*) tumor suppressor gene, which results in the uncontrolled activation of hypoxia-inducible transcription factors (HIFs)^3–6^. Accordingly, therapies aimed at targeting hypoxia signaling, along with immunotherapies, have significantly advanced the treatment of ccRCC^7,8^. However, these treatments are generally less effective against other forms of kidney cancer, which are driven by different biological mechanisms^9,10^.

Translocation RCC (tRCC) is a rare and aggressive subtype of RCC defined by oncogenic gene fusions involving a transcription factor in the MiT/TFE family, usually TFE3^11–13^. tRCC comprises approximately 1-5% of RCCs in adults and about 50% of RCCs in children; however, its true incidence may be underestimated due to frequent histologic misclassification as other, more common, subtypes of kidney cancer^14–16^. There is no established standard of care treatment specifically for tRCC, and responses to immune checkpoint inhibitors (ICI) and vascular endothelial growth factor (VEGF)-targeted kinase inhibitors are typically more modest than in ccRCC^17–20^. Moreover, an incomplete molecular understanding of tRCC has hindered the ability to potentiate these therapies and to identify new mechanism-informed targets in this subtype.

Prior molecular characterization studies of tRCC have been limited in scale due to its rarity. Nonetheless, recent whole exome sequencing (WES) and whole genome sequencing studies have revealed that tRCCs have a low mutational rate and harbor few recurrent genomic alterations apart from the driver MiT/TFE gene fusion^12,21–26^. Despite this quiet and homogeneous genomic profile, tRCCs have been proposed to comprise multiple transcriptional and proteomic subtypes, though these studies have relied on bulk profiling and have identified different sets of conserved molecular programs^22,23^. Indeed, multiple pathways have been identified as activated in tRCC, including the antioxidant response, mTOR signaling, oxidative phosphorylation, autophagy, epithelial-mesenchymal transition, Ret signaling and Wnt signaling, amongst others^12,22,23,27–31^.

To date, malignant programs in tRCC have not been systematically studied at single-cell resolution, nor have they been linked to specific components of the tumor ecosystem in tRCC. Therefore, how oncogenic TFE3 fusions direct malignant programs, shape the immune landscape, and influence therapy response in tRCC remains poorly defined. In this study, we assembled a cohort of 20 tRCC samples which were profiled using single-nucleus RNA sequencing (snRNA-seq), along with single nucleus ATAC sequencing (snATAC-seq), spatial transcriptomic analysis, and T Cell Receptor (TCR) sequencing. By integrating 128 bulk transcriptomic tRCC profiles from prior studies^22,23,32–34^ and a systematic comparison with 1300 published ccRCC bulk and 32 single-cell transcriptional profiles^32,33,35^, we present a detailed picture of how TFE3 fusions drive malignancy and contribute to immunotherapy resistance, while nominating new therapeutic entry points in this aggressive cancer.

## RESULTS

### Single nucleus sequencing of tRCC

We assembled a cohort of 20 tRCC samples from 15 patients, of which 17 samples (16 tumors, 1 adjacent normal) underwent single-nucleus profiling via one of three platforms: **(1)** 10x Genomics 3’ snRNA-Seq (6 fresh frozen samples); **(2)** 10X 3’ multiome (5 fresh frozen samples)**; (3)** 10x-Flex fixed RNA-Seq (6 FFPE samples). (**Figure 1A & S1A**). For comparative analysis, we uniformly reprocessed data from 32 snRNA-seq and 24 snATAC-seq samples of ccRCC profiled via the 3’ multiome platform in a prior study^35^. We also aggregated 128 tRCC and 1311 ccRCC bulk transcriptomes from across 5 published studies for validation analyses^22,23,32–34^ (**Figure 1A**).

**Figure 1.**
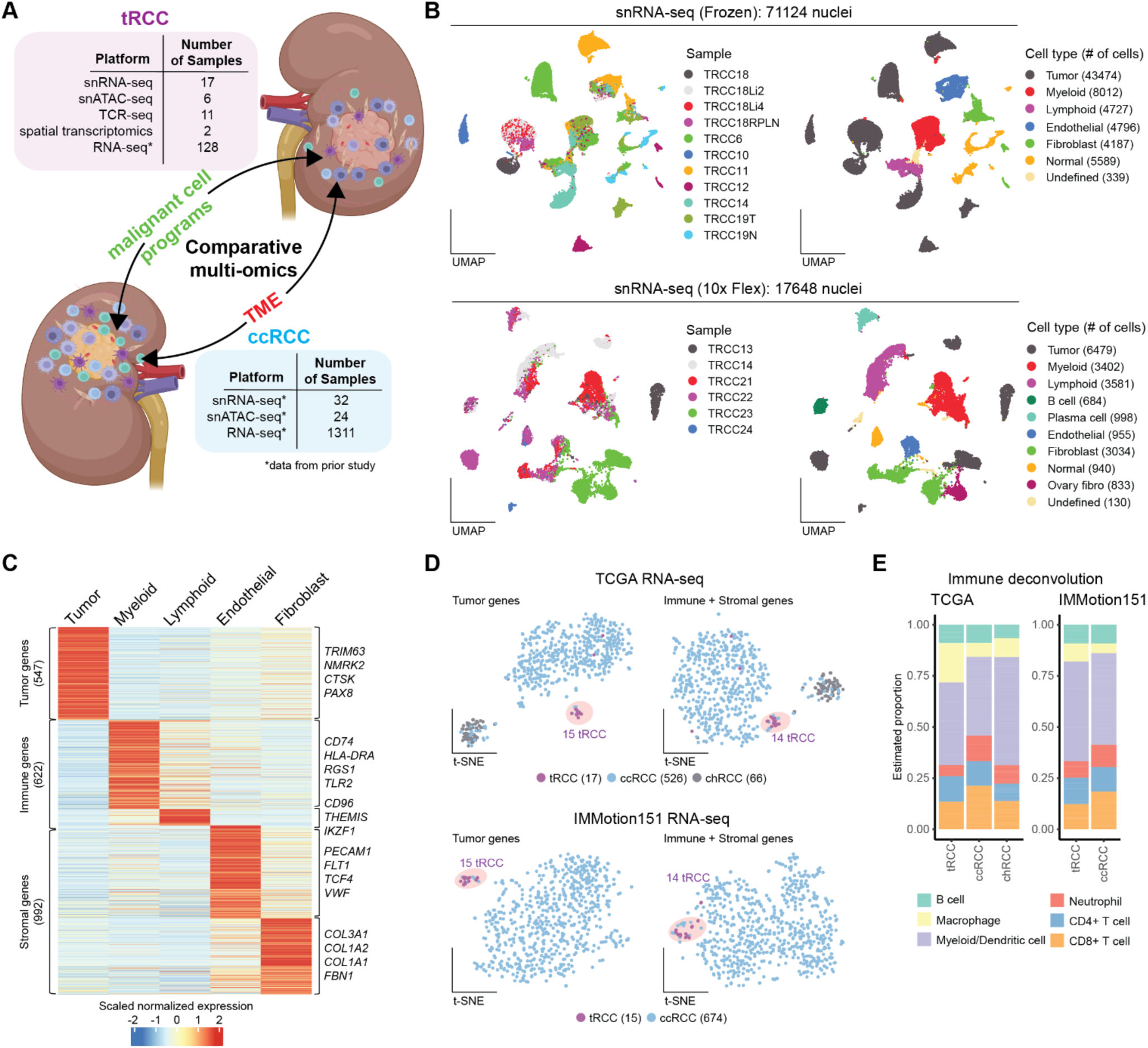
tRCCs and ccRCCs have distinct cellular landscapes. A. Schematic representation of study design, including sample numbers for tRCC and ccRCC cohorts analyzed in the study. B. Uniform manifold projections (UMAP) of nuclei from tRCC samples annotated by Sample ID (left) and cell lineage (right) in snRNA-seq frozen (top) or 10x-flex (bottom) cohorts. C. Heatmap of genes highly expressed in tumor, immune, or stromal cells in tRCC based on snRNA-seq data (frozen cohort). D. Projection of genes from (C) onto bulk RNA-Seq RCC datasets (top, TCGA; bottom, IMmotion151), with t-SNE plots showing separation of subtypes based on either tumor (left) or microenvironmental (immune + stromal, right) gene signatures. E. Stacked bar plots displaying estimated proportions of immune cell types in various RCC subtypes, derived from immune deconvolution analysis of bulk RNA-Seq datasets.

Our snRNA-Seq cohort was female predominant (8 females, 4 males), consistent with the known female sex-bias in tRCC^21^. Four distinct *TFE3* fusion partner genes were represented (*PRCC*, *SFPQ*, *MED15*, and *LUC7L3*). Given the rarity of tRCC, the cohort was heterogeneous with respect to tumor site (11 primary, 5 metastases) and treatment history (8 treatment naïve, 8 with prior systemic therapy) (**Figure S1A & Table S1**).

Unsupervised clustering of snRNA-seq data from our tRCC cohort revealed robust separation of non-malignant cell types such as endothelial cells, fibroblasts, immune populations, and normal kidney cells, regardless of patient origin or platform. By contrast, malignant cells segregated by patient, suggesting a degree of intertumoral heterogeneity (**Figure 1B & S1B-C**). While frozen and FFPE snRNA-seq platforms yielded relatively comparable feature distributions, cell type proportions differed between platforms (likely due to differences in sample preparation), leading us to analyze the two cohorts separately (**Figure S1D-E**).

### tRCC exhibits distinctive tumor and microenvironment-derived gene expression

After assigning cell types in our tRCC snRNA-seq cohort, we identified genes differentially expressed in the tumor, immune, and stromal compartments (**Figure 1C**). We then mapped these lineage-specific signatures onto two bulk RNA-seq datasets that included tRCCs and other kidney cancers (TCGA; IMmotion151)^32,33^. tRCC transcriptomes segregated from both ccRCC and chromophobe RCC (chRCC) when clustered based on tumor genes (**Figure 1D**, left) and when clustered by immune/stromal genes (**Figure 1D**, right) (Note: papillary RCCs were excluded from both cohorts due to their molecularly heterogeneous nature^36,37^). Immune deconvolution analyses (CIBERSORTx) inferred increased tumor-associated macrophages (TAM) in tRCC compared to both ccRCC and chRCC, as well as reduced CD8+ T cells in tRCC relative to ccRCC (**Figure 1E & S1F**). These analyses suggest that the unique clinical features of tRCC may be linked to distinctions both in malignant programs as well as in the surrounding stromal and immune microenvironment.

### Divergent transcriptional and epigenetic programs in tRCC and ccRCC

To characterize the malignant programs that typify tRCC, we performed differential gene expression analysis on malignant cells from our tRCC snRNA-seq cohort (N=10 frozen samples only) and a previously published snRNA-seq cohort of ccRCC (N=32)^35^. We intersected tRCC-enriched genes from this analysis with differential gene expression performed on a much larger bulk RNA-seq cohorts for tRCC (N=95^22,32,33^) and ccRCC (N=1311^32,33^) (**Figure 2A & Table S2**). This consensus set of genes upregulated in tRCC tumors was strongly enriched for pathways associated with reactive oxygen species (ROS) handling, oxidative phosphorylation, and the electron transport chain, consistent with our recent finding that these pathways are under direct transcriptional control of the TFE3 fusion^27^. tRCCs also exhibited an upregulation of the lysosomal pathway, consistent with the canonical role for MiT/TFE genes as master regulators of lysosomal biogenesis^38^ (**Figure 2A**). By contrast, ccRCC tumors displayed upregulation of hypoxia and glycolysis related genes, in keeping with the well-established role of HIF stabilization and Warburg-like metabolism in ccRCC^5,39^ (**Figure S2A**).

**Figure 2.**
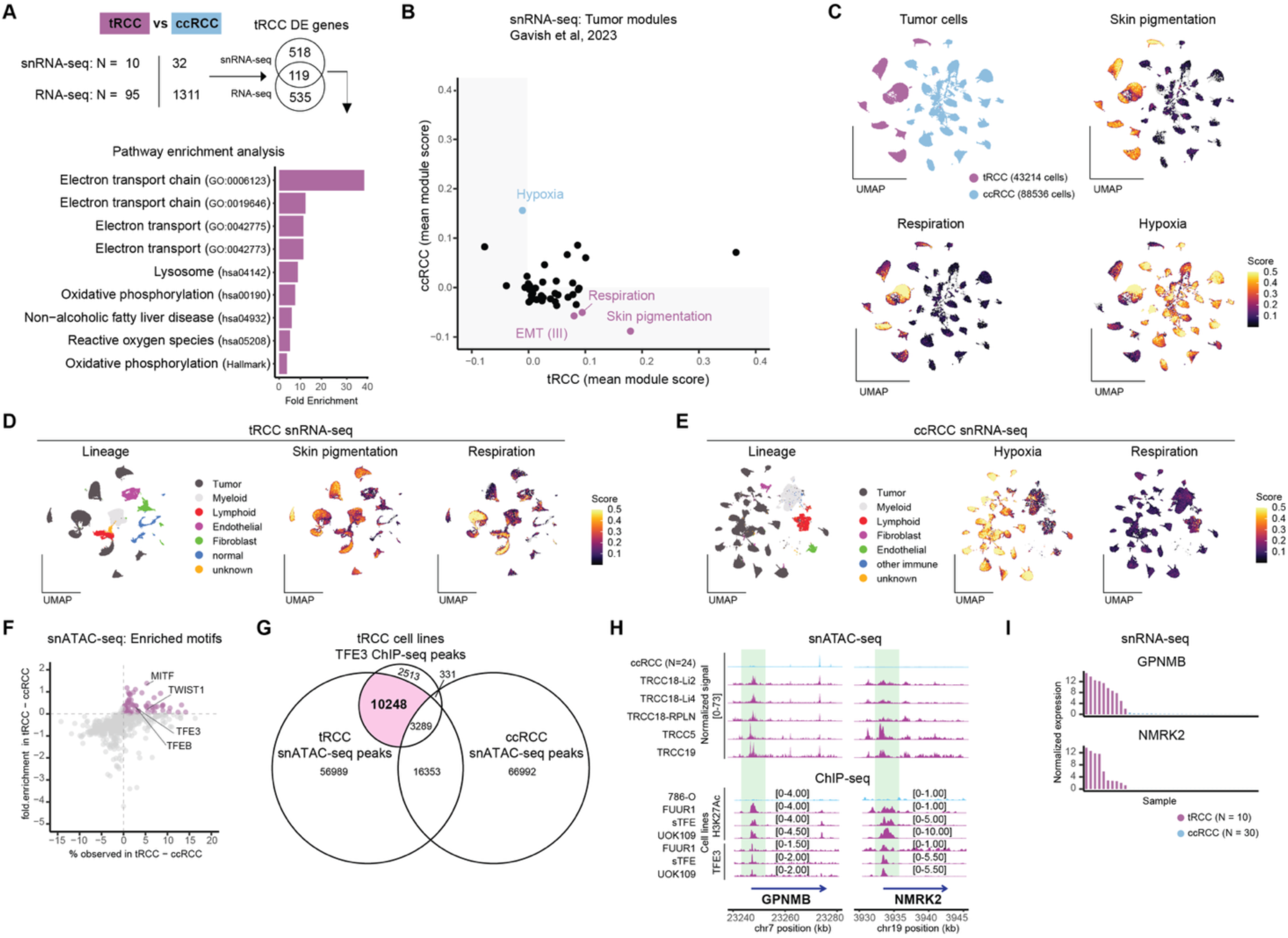
Divergent tumor transcriptional profiles in tRCC and ccRCC. A. Differential expression analysis of tRCC vs ccRCC tumors using snRNA-seq (Frozen) and RNA-seq data, highlighting top enriched pathways in tRCC tumors identified based on gene-ontology. B. Scatterplot of program enrichment in tRCC vs ccRCC tumor cells from snRNA-seq data (Frozen), based on consensus meta-programs established in a recent pan-cancer study^40^. C. UMAPs representing tumor cells from tRCC and ccRCC datasets (top left), colored by: skin pigmentation program module score (top right), respiration program module score (bottom left), and hypoxia program module score (bottom right). D. UMAPs showing all lineages in the tRCC snRNA-seq dataset (left) colored by: skin pigmentation module score (middle) and respiration module score (right). E. UMAPs depicting all cellular lineages in the ccRCC snRNA-seq dataset (left) colored by: hypoxia module score (middle) and respiration module score (right). F. Scatterplot of motifs enriched in tRCC and ccRCC tumor cells from snATAC-seq data. X-axis represents the difference in percent motif observed in tRCC vs. ccRCC tumor cells and Y-axis represents the difference in fold enrichment of each motif in tRCC vs. ccRCC. tRCC-enriched motifs are highlighted in pink. G. Venn diagram showing the intersection of peaks from tRCC snATAC-seq data (tumor cells), TFE3 ChIP Seq from tRCC cell lines, snATAC-seq from ccRCC tumor cells. Peaks specific to tRCC are highlighted in pink. H. Peaks at the loci for two tRCC-marker genes (*GPNMB* and *NMRK2*) from: snATAC-seq of tRCC and ccRCC tumors (top) and ChIP-seq of H3K27ac or TFE3 in tRCC (FUUR1, sTFE, UOK109) or ccRCC (786-O) cell lines (bottom). I. Waterfall plot representing normalized expression of tRCC-specific genes, *GPNMB* and *NMRK2*, in each sample of tRCC and ccRCC tumors profiled via snRNA-seq.

We also evaluated cancer cell states previously defined in pan-cancer single cell studies^40,41^. Consistent with our above analyses and recent work suggesting metabolically divergent states in tRCC and ccRCC^27,28^, a respiration program was enriched in tRCC while a hypoxia program was enriched in ccRCC (**Figures 2B-E & S2B**). tRCC malignant cells also displayed enrichment of epithelial-mesenchymal transition (EMT) and skin pigmentation transcriptional programs, the latter being originally described as an MITF-driven hallmark module in melanoma^40^ and reflective of tRCC being driven by an MiT/TFE family transcription factor (**Figure 2B-C**). Both the pigmentation and respiration programs were also enriched in tumor cells relative to non-malignant cells, with the exception of some activity of the pigmentation program in myeloid cells (consistent with a role for TFE3 in macrophages^42–44^) and some activity of the respiration program in lymphoid cells (consistent with resting T lymphocytes preferring aerobic respiration^45^) (**Figure 2D**).

To assess whether these transcriptional differences were driven by distinct chromatin accessibility profiles in tRCC vs. ccRCC malignant cells, we interrogated our tRCC single snATAC-seq dataset (23542 nuclei from 5 samples) dataset together with a published ccRCC snATAC-Seq dataset (119130 nuclei from 24 samples^35^). As expected, transcription factor motif enrichment analysis identified increased accessibility of MiT/TFE family members in tRCC vs. ccRCC (**Figure 2F**). We also observed tRCC-specific enrichment of a TWIST1 motif (master regulator of EMT^46^), consistent with our observation of an EMT transcriptional program in tRCC malignant cells.

To determine whether regions of chromatin accessibility enriched in tRCC marked fusion target genes, we next intersected TFE3 fusion binding sites as determined by ChIP-Seq on tRCC cell lines^27^ with snATAC-seq peaks in the tRCC and ccRCC cohorts. A majority of TFE3 fusion ChIP-Seq peaks overlapped with tRCC-specific ATAC-Seq peaks (10248 peaks, hypergeometric test p = 6e-85) (**Figure 2G**). These regions exhibited increased H3K27Ac signal and TFE3 occupancy of the canonical TFE3 target genes, *GPNMB*^30^ and *NMRK2*^47,48^ (**Figure 2H**). Correspondingly, these fusion target genes were also strongly upregulated at the RNA level in tRCC tumors (**Figure 2I**), highlighting the direct linkage between chromatin accessibility and gene expression downstream of TFE3 fusions.

### Cell-type identification points to a common cell of origin for tRCC and ccRCC

The stark transcriptional and epigenetic differences between tRCC and ccRCC malignant cells may reflect their distinct genetic drivers or a distinct cellular origin, or both. Kidney cancers are known to be cellularly diverse, comprising dozens of histologies originating from various cell types along the length of the nephron^49^. For example, the cell of origin for ccRCC has been proposed as the proximal tubule, while collecting duct intercalated cells are proposed as the cell of origin for chromophobe RCC, and proximal tubule for papillary RCC^49–51^. Recent studies have proposed a proximal tubule cell origin for tRCC^51,52^; however, owing to the rarity of tRCC, these conclusions were based on bulk transcriptome analysis or on single-cell profiling of a single sample and not mapped to the resolution of a subpopulation within proximal tubule cells. Therefore, we sought to comprehensively define the cell of origin in tRCC.

We leveraged a recently published snRNA-seq atlas of normal adult human kidney^53^ to classify cell types within a tumor-adjacent normal sample from patient TRCC19, identifying 13 distinct cell types representing all segments of the nephron (**Figure S2C-E).** We then quantified the transcriptomic similarity between malignant cells in each tRCC tumor and each normal reference cell type. Across tRCC malignant cells, we found a substantial similarity to the vascular cell adhesion molecule-1 (*VCAM1*)-positive proximal tubule cells (PT_VCAM1) (**Figure S2F-G**). PT_VCAM1 cells represent a subpopulation of proximal tubule cells that also show increased expression of kidney injury molecule-1 (KIM1, encoded by *HAVCR1*) and may represent cells with progenitor-like features that are expanded in the setting of renal injury^53–55^. Of note, plasma KIM1 has emerged as a biomarker of malignant pathology for kidney masses, recurrent RCC, and as a predictive marker of response to immunotherapy in metastatic RCC^56,57^. Notably, while malignant cells in some tRCC samples (e.g. TRCC6, TRCC11, TRCC19) showed highest similarity to parietal epithelial cells (PEC), which line Bowman’s capsule, *VCAM1* has also been shown to be expressed in these cells, as well as in scattered other cell types throughout the kidney^53,58^ (**Figure S2E, G)**. A parallel analysis on ccRCC tumors^35^ showed uniform mapping of ccRCC malignant cells to the same PT_VCAM1 subpopulation, consistent with a prior report implicating a *VCAM1*-expressing proximal tubule 1 (PT1) cell as the likely cell of origin for ccRCC^50^ (**Figure S2H**). These findings were further supported by analysis of bulk RNA-seq datasets, which similarly suggested proximal tubule origin for both tRCC and ccRCC^49,51,52^ (**Figure S2I)**. Altogether, these findings suggest that tRCC and ccRCC most often arise from a shared *VCAM1*-expressing proximal tubular cell of origin but undergo divergent transcriptional and epigenetic reprogramming in the context of distinct oncogenic drivers.

### Conserved oncogenic meta-programs in malignant tRCC cells

We next sought to identify gene expression programs that were shared across multiple tumors in our cohort. We identified co-regulated gene modules via non-negative matrix factorization (NMF), converging upon six tumor meta-programs (MPs) that were each defined by the expression of canonical markers (**Figure 3A & Table S3**): **(MP1)** Cell cycle (*CDK1*, *TOP2A*, *MI67*); **(MP2)** EMT (*COL1A1*, *VCAN, FN1*); **(MP3)** Angiogenesis (*VEGFA*, *SERPINE1*); **(MP4)** Inflammation (*CXCL10*, *IFIT1*); **(MP5)** TNFα (*FOS*, *JUN*); **(MP6)** Proximal tubule (PT) (*HNF4A*, *SLC13A1*).

**Figure 3.**
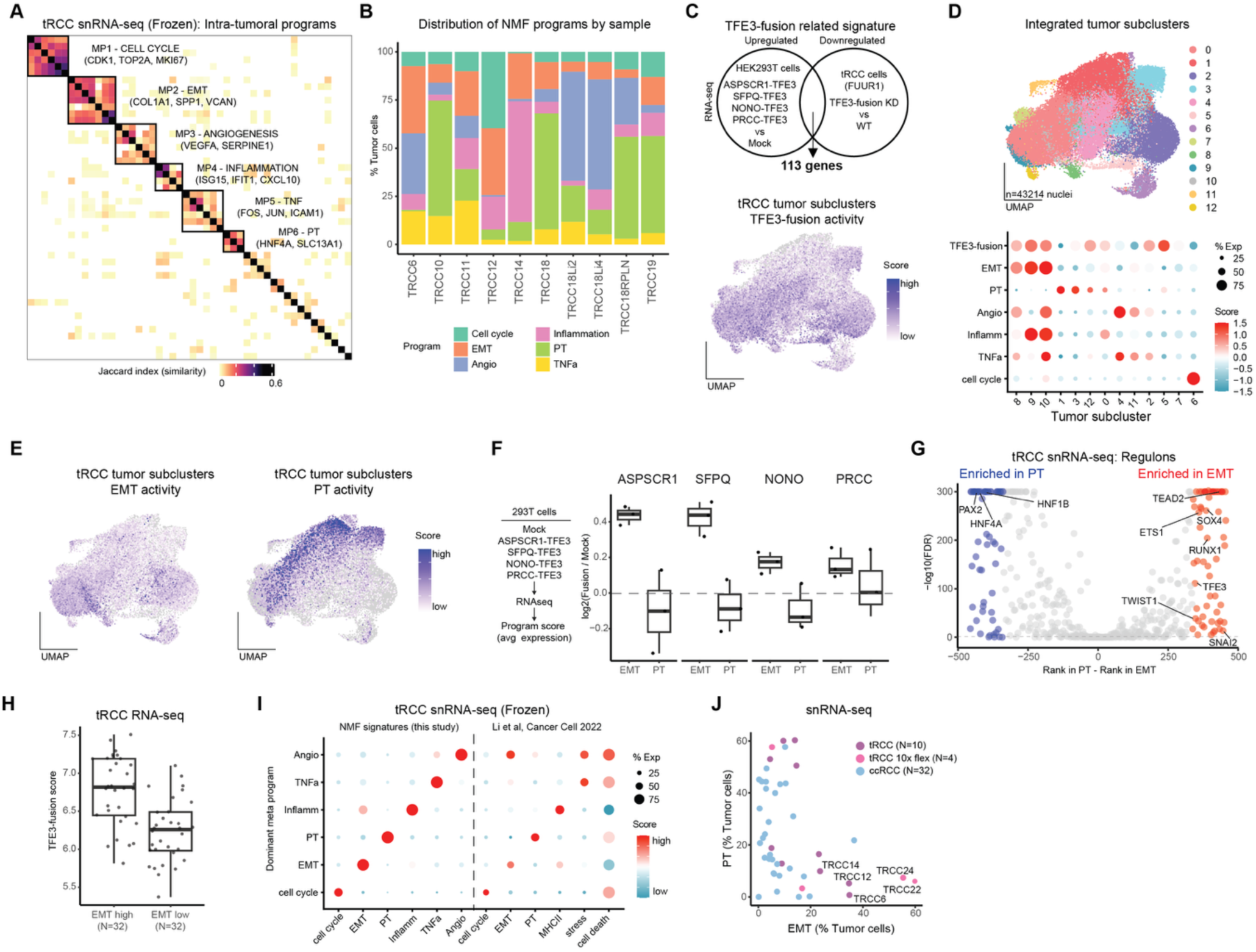
Consensus tumor meta-programs in tRCC. A. Heatmap indicating similarity (Jaccard index) between individual tumor NMF programs in tRCC frozen cohort, with six conserved tRCC meta-programs (MP1-MP6) annotated. B. Stacked bar plot showing the distribution of identified NMF programs across the tRCC snRNA-seq (frozen) cohort. C. TFE3-regulated genes identified in *in vitro* gain- and loss-of-function experiments^12,27^ (top) were used to derive a TFE3 fusion activity signature. Bottom, UMAP representing TFE3-fusion activity score across tRCC tumor cells (frozen cohort). D. UMAP representing subclusters from integrated tRCC tumor cells (top) and dot plot showing relative scores of tumor NMF programs as well as TFE3 fusion activity signature in each tumor subcluster in frozen cohort (bottom). E. UMAPs representing EMT activity score (left) and PT activity score (right) in tRCC tumor cells (frozen cohort). F. Box plots depicting changes in EMT and PT scores upon overexpression of various different TFE3-fusions in HEK-293T cells^12^. G. Volcano plot representing transcription factor regulons identified in EMT vs PT cells from tRCC tumor snRNA-seq data. X-axis represents the difference in ranks of regulons in PT vs EMT cells. Regulons enriched in EMT are highlighted in red while those enriched in PT are highlighted in blue. H. Boxplots showing TFE3-fusion activity score in EMT-high vs EMT-low tumors from bulk RNA-Seq data of tRCC. I. Dot plot showing association between tRCC NMF programs identified in this study (frozen cohort) and NMF programs defined previously in ccRCC^63^. J. Scatterplot showing proportion of EMT and PT tumor cells in each tRCC and ccRCC sample.

All six MPs were expressed to varying degrees in malignant cells in each tumor, indicative of substantial transcriptional heterogeneity, despite tRCC being a genetically quiet tumor with few recurrent genomic alterations apart from the driver fusion^12,21–24,26^. The PT MP was the most prevalent across all malignant cells (23.8% of all malignant cells overall; range 0.7%-60.2% per sample), reinforcing a proximal tubular epithelial cell origin for tRCC. The cell cycle MP, likely reflective of rapidly cycling cells, was mutually exclusive with all other MPs; a recent scRNA-seq study has suggested enrichment of a similar program amongst tumor progenitor cells in synovial sarcoma^59^ (**Figure 3B-D & S3A**).

Intriguingly, ∼20% of malignant cells (1%-57%) in tRCC expressed an angiogenesis MP, despite the fact that tRCCs do not harbor *VHL* loss and therefore do not have genetically enforced activation of angiogenesis via HIF-2a signaling, as in ccRCC^3–6^. Nonetheless, the expression of an angiogenesis MP in a subset of malignant tRCC cells is consistent with the clinical observation of improved outcomes with VEGF targeted therapy or immunotherapy-VEGF combinations (IO/TKI) rather than immunotherapy alone in tRCC^17,60^. Notably, the angiogenesis MP was frequently co-expressed with either the inflammation or TNFa MPs, or both, perhaps suggesting activation of angiogenesis by inflammatory cytokines in tRCC^61,62^ (**Figure 3D**).

The EMT and PT MPs were mutually exclusive in malignant cells, consistent with a recent scRNA-seq study of ccRCC^63^. This held across patients, with malignant cells across the entire cohort integrated into 12 subclusters (**Figure 3D-E**). Moreover, snRNA-seq of the primary tumor and three metastatic sites from a recently reported tRCC rapid autopsy case^21^ revealed that The EMT MP was stronger in the metastatic sites relative to the primary tumor, while the PT MP showed the inverse pattern (**Figure S3B-D**). These findings demonstrate, from malignant cells within the same patient, a canonical association of the EMT with metastatic colonization^64^.

We next sought to determine to what extent these MPs were directly linked to TFE3 fusion activity. We derived a consensus signature of 113 fusion-regulated genes based on RNA-seq data from two *in vitro* perturbation experiments: overexpression of TFE3 fusion constructs in HEK293T cells^12^ and knockdown of the endogenous *TFE3* fusion in tRCC cell line, FUUR1 (**Figure 3C & Table S3**)^27^. Malignant cells displayed substantial heterogeneity in the fusion transcriptional signature score, with a higher signature score associated with increased utilization of the EMT MP and inversely associated with utilization of the PT MP (**Figure 3C-E**). This relationship was recapitulated *in vitro* by overexpression of four different *TFE3* fusions in HEK293T embryonic kidney cells^12^ (**Figure 3F**).

In further support of TFE3 activity skewing towards an EMT cell state, regulon analysis revealed that EMT-high malignant cells were enriched for motifs associated with TFE3 along with canonical EMT-associated transcription factors including TWIST1, SOX4, RUNX1, SNAI2, ETS1, and TEAD2^46,65–69^; conversely, lineage factors associated with renal epithelial cell identity (e.g. PAX2, HNF4A, and HNF1B^70–72^) were enriched in PT-high cells (**Figure 3G**). We also analyzed bulk RNA-Seq data from 128 tRCC tumors across 5 published cohorts^22,23,32–34^ and confirmed that a higher TFE3 fusion signature score was positively correlated with an EMT signature score and inversely correlated with a PT signature score (**Figure 3H & S3E**).

Altogether, our observation of a PT MP conserved across all samples supports a proximal tubule cell of origin for tRCC, while its mutually exclusive expression with an EMT program is suggestive of a fusion-driven mesenchymal differentiation program leading to loss of epithelial cellular identity.

### Comparative analysis of meta-programs in ccRCC

As tRCC is a rare and biologically unique tumor type not well-represented in recent genomic atlas efforts, we next sought to contextualize our conserved tRCC MPs with respect to those identified in other cancers. We first compared our tRCC oncogenic MPs with those recently defined in a cohort of 10 ccRCC samples profiled by scRNA-seq^63^. All six tRCC MPs overlapped substantially with one or more gene programs identified in ccRCC: EMT, cell cycle, and PT programs were identified in both studies while our tRCC MPs of inflammation and TNFa mapped remarkably well to MHCII and stress/cell death programs identified in ccRCC, respectively; the tRCC angiogenesis MP mapped variably to the EMT, stress, and cell death MPs in ccRCC (**Figure 3I**). Despite several malignant cell programs being shared between tRCC and ccRCC, there were stark differences in their utilization: most notably, tRCC samples more commonly showed substantial EMT MP utilization relative to ccRCC (6/14; 43% for tRCC vs. 1/32; 3% samples for ccRCC at threshold of >20% tumor cells) (**Figure 3J & S3F**).

Our tRCC MPs also mapped to states identified in a recent comprehensive pan-cancer single cell study of >1100 tumors spanning 24 tumor types^40^ (**Figure S3G**). The tRCC EMT MP mapped strongly to several EMT states identified in this pan-cancer study, as well as to a mesenchymal program (MES) identified in glioma. Interestingly, the tRCC EMT MP also overlapped with a skin pigmentation program associated with MITF activity in melanoma – consistent with our results above that implicate the TFE3 fusion (an MITF homolog) in directly driving EMT. Our tRCC PT MP exhibited a strong correlation with a pan-cancer glutathione program enriched in ccRCC, and consistent with a critical role for glutathione handling in renal proximal tubule cells^73^. The tRCC angiogenesis MP matched to the pan-cancer hypoxia program but also had partial alignment with many additional programs; together with its variable matching against ccRCC MPs, as discussed above, this suggests that this particular MP may be somewhat unique to tRCC, although we cannot exclude signature-related differences due to sample preparation or platform.

Overall, our results suggest that, despite harboring a dominant genetic driver distinct from almost all other cancers, tRCC cells converge on broadly conserved core oncogenic programs observed in other solid tumors.

### Dysfunctional T cell phenotypes and response to ICI in tRCC

Responses to immunotherapy vary widely across kidney cancer subtypes: while ccRCC is generally considered an immune-responsive tumor, response to immunotherapy in other RCCs is more variable, with extremely low immune-responsiveness in certain subtypes such as renal medullary carcinoma (RMC)^74^ and chromophobe RCC (chRCC)^75^. Several small studies suggest some tRCCs do respond to immunotherapy, albeit at a lower rate than ccRCC^12,17,22,76^. Variable immune-responsiveness across RCC subtypes likely stems both from differences in the quantity and nature of tumor antigens capable of stimulating an anti-tumor immune response, and in microenvironmental factors that may constrain anti-tumor immunity.

We therefore sought to define the tumor microenvironment (TME) composition in tRCC, particularly in light of our initial observation that tRCCs diverge from ccRCCs on the basis of immune/stromal gene expression programs (**Figure 1E**). Focusing initially on lymphocytes, cell type identification revealed CD4⁺ T cells, CD8⁺ T cells, regulatory T cells (Tregs), natural killer (NK) cells, B cells, and plasma cells in tRCC tumor samples (**Figure 4A**; 10x-Flex fixed RNA-Seq cohort was used for this analysis due to greater representation of lymphocytes, **Figure S1E**).

**Figure 4.**
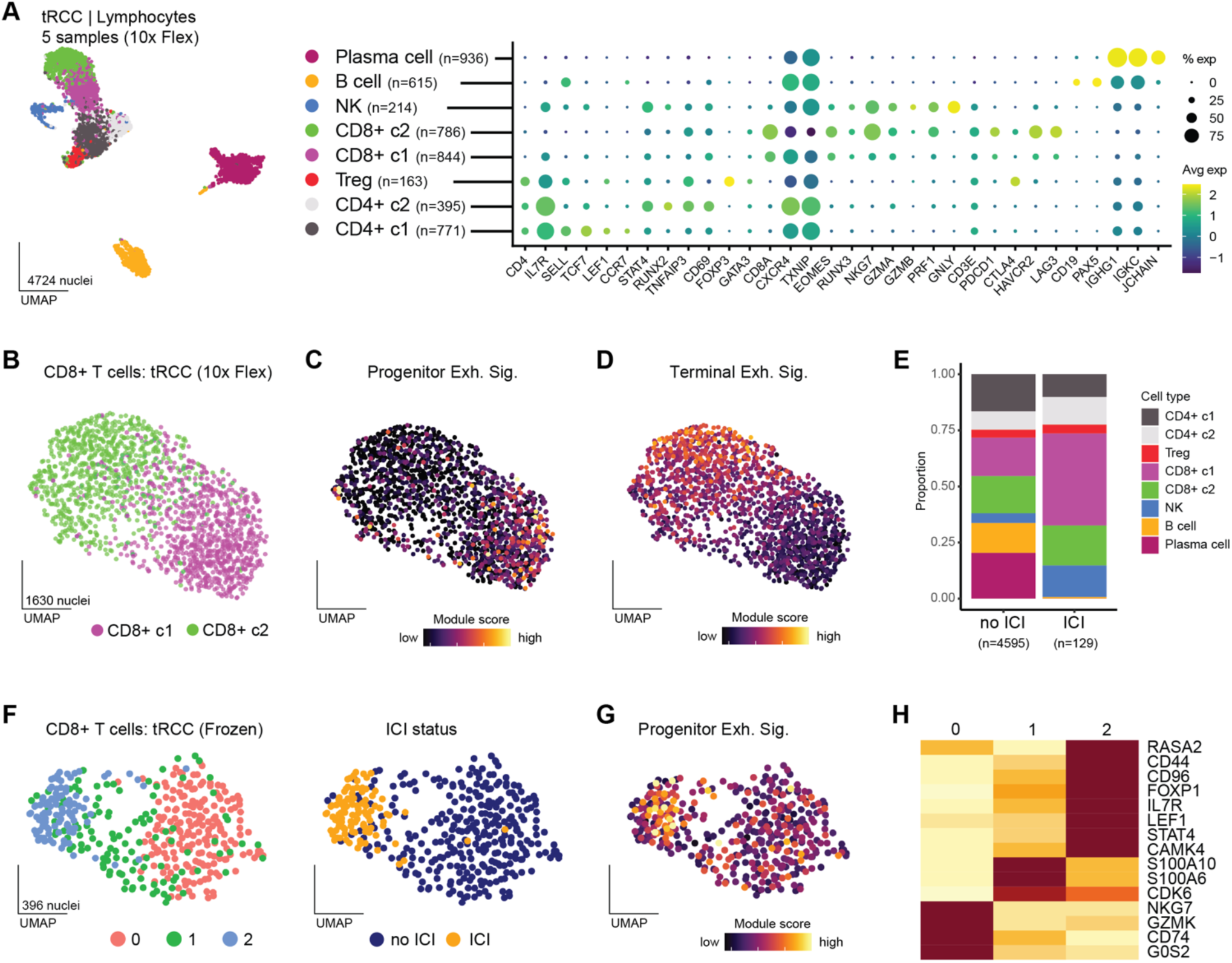
CD8 T cell states in the tRCC TME. A. UMAP representation of all lymphocytes (left) as well as annotation and marker genes of lymphocyte subsets (right) in the tRCC (10x Flex) cohort. B. UMAP of CD8 T cell clusters (10x Flex cohort). C. UMAP representing progenitor exhausted signature score across CD8 T cells (10x Flex cohort). D. UMAP representing terminally exhausted signature score across CD8 T cells (10x Flex cohort). E. Stacked bar plot depicting proportion of lymphocyte subtypes in tRCC samples from ICI-naïve (left) and ICI-treated (right) patients. F. UMAP of CD8 T cell clusters (left) colored by ICI status (right) in the tRCC frozen cohort. G. UMAP representing progenitor exhausted signature score across CD8 T cells from the tRCC frozen cohort. H. Heatmap with top expressed marker genes associated with CD8 T cell subclusters from the tRCC frozen cohort.

Notably, despite an overall decreased T lymphocyte abundance compared with ccRCC (**Figure 1E** & ^12^), both CD4+ and CD8+ T cells were still present in the tRCC TME (**Figure S4A**). This finding is a notable contrast from chRCC, which is a typically “cold” tumor with an exceedingly low proportion of infiltrating immune cells^75,77^. Among CD4+ T cells, we observed two clusters: CD4 c1, with expression of *IL7R*, *TCF7*, and *CCR7* consistent with a naïve/central memory phenotype; and CD4 c2, which exhibits features of activated/effector memory phenotype with expression of *STAT4*, *RUNX2*, *CD69*, and *TNFAIP3*^78,79^. Together with the presence of abundant plasma cells (**Figure 4A**), the presence of central memory-type CD4 cells may be suggestive of the presence of an immature tertiary lymphoid structures (TLS) – ectopic lymphoid formations resembling secondary lymphoid organs – which, in their mature form, have recently been linked to immune-responsiveness in ccRCC^80,81^. Additionally, although tRCC tumors displayed sparse NK cell infiltrate, these NK cells skewed toward cytotoxicity rather than a dysfunctional tumor-resident phenotype that has been recently linked to poor outcomes in ccRCC^82^ (**Figure S4C**).

Given our finding that tRCCs do indeed possess CD8+ tumor-infiltrating lymphocytes (TILs), we next turned our attention to defining CD8+ T cell states that might modulate immune responsiveness. Prior studies have elucidated heterogeneity in exhausted CD8+ TIL populations: progenitor-exhausted CD8+ TILs exhibit polyfunctionality, persistence, share properties with exhausted CD8+ T cells in chronic viral infection models and are capable of being reinvigorated by ICI, while terminally-exhausted CD8+ T cells display greater cytotoxicity but have reduced long-term survival^83^. Reclustering of CD8+ cells in our 10X flex cohort revealed two subclusters (**Figure 4B**): CD8 c1 was enriched for markers of progenitor-exhausted T cells, such as *IL7R*, *TCF7*, and *CXCR4*, while CD8 c2 expressed markers more typical of terminal exhaustion including *PDCD1*, *LAG3*, and *HAVCR2*^78,83,84^ (**Figure 4C-D**). Moreover, progenitor-exhausted CD8+ T cells were significantly more abundant in tumors from patients who had been exposed to ICI, consistent with transfer experiments in mouse models, which suggest that ICI acts upon the progenitor-exhausted CD8+ TIL population^83^ (**Figure 4E**). These results were confirmed in the frozen snRNA-seq sub-cohort, where clustering of CD8+ T cells revealed three clusters, with clusters 0 and 1 being primarily derived from ICI-naive tumors and cluster 2 enriched in ICI-treated samples and displaying a higher progenitor-exhausted signature (**Figure 4F-G**). Functional annotation of these clusters via marker gene expression and gene signature scoring revealed a progression from naïve/quiescent (cluster 0; *G0S2*, *CD74*^85,86^) to proliferative/activated (cluster 1; *CDK6*, *S100A6*, *S100A10*), and ultimately to a progenitor-exhausted/memory-like phenotype (cluster 2; *IL7R*, *FOXP1*, *CD44*, *CD96*, *LEF1*, and *RASA2*^87–92^) (**Figure 4G-H**). Metabolic pathway module scores were also consistent with a naïve-like phenotype in cluster 0 (high oxidative phosphorylation^93^), a proliferative phenotype in cluster 1 (high glycolysis/mTOR signaling^94^), and a memory-like phenotype in cluster 2 (high fatty acid oxidation^93^) (**Figure S4B**).

Although tRCC has a very low mutational burden^12^, the driver fusion may represent a strong and clonal neoantigen, given that the *TFE3* fusion junction spans two genes that are not normally juxtaposed^26^. To evaluate for clonal expansion of tumor-responsive T cells, we performed TCR sequencing on 11 tRCC tumor samples (of which 8 tumors were in our single-cell cohort). After excluding publicly reported human viral antigen T cell receptor sequences, we observed a median of 485 TCR-α and 890 TCR-β non-viral unique clonotypes per sample (range 345 to 2535 TCR-α & 519 to 4208 TCR-β), which were classified as hyperexpanded (> 3%), large (> 1%), medium (> 0.01%), and small (< 0.01%) groups based on their frequency in each sample (**Figure S4D-F**). Sample TRCC14 sample was an outlier in terms of number of unique clonotypes; this sample also had the highest lymphocyte infiltration and interferon module score on snRNA-seq profiling (**Figure S4E**). However, all samples had large or hyperexpanded TCR clones to varying degrees, suggesting the presence of expanded TIL populations responding to tumor-specific antigens. Nonetheless, due to the range of potential *TFE3* fusion partners and breakpoints^21^ coupled with HLA-type diversity, the strength and quality of a fusion-derived neoantigen may vary considerably between patients.

Overall, our analyses point to progenitor-exhausted CD8+ T cells and clonally-expanded TILs in tRCC – including after ICI exposure – suggestive of capacity for an effective anti-tumor T cell response. However, despite enrichment of progenitor-exhausted CD8+ T cells following ICI exposure, terminally exhausted CD8+ TILs were not proportionally expanded (**Figure 4E**). This implies an immunologically active yet functionally restrained TME in tRCC, which may explain the limited clinical efficacy of single agent PD1/PD-L1 inhibitors in tRCC.

### Suppressive macrophages and fibroblasts in the tRCC TME

We next sought to determine if the constitution of the myeloid compartment in tRCC could reconcile the presence of tumor-specific TILs capable of responding to ICI with the typically modest responses to ICI clinically observed in tRCC^17,95^. By proportion, myeloid cells constituted the largest compartment of the tRCC TME (**Figure S1E**), which included a number of different macrophage subtypes, including: SPP1+ macrophages, C1Q+ macrophages, tissue resident macrophages (Mac-TRM), and GPNMB macrophages (**Figure 5A & S5A-5B**). Comparison of macrophage subtypes in a published ccRCC snRNA-Seq dataset^35^ revealed stark differences with tRCC. SPP1+ macrophages, recently implicated in the suppression of CD8+ TIL activity and adverse clinical outcomes across several tumor types^96–98^, comprised approximately 25% of macrophages in tRCC but were under represented in the ccRCC cohort. By contrast, IFN-primed macrophages, which are associated with anti-tumor effects^99,100^, constituted ∼10% of the myeloid population in ccRCC but were undetectable in tRCC (**Figure 5B & S5C**).

**Figure 5.**
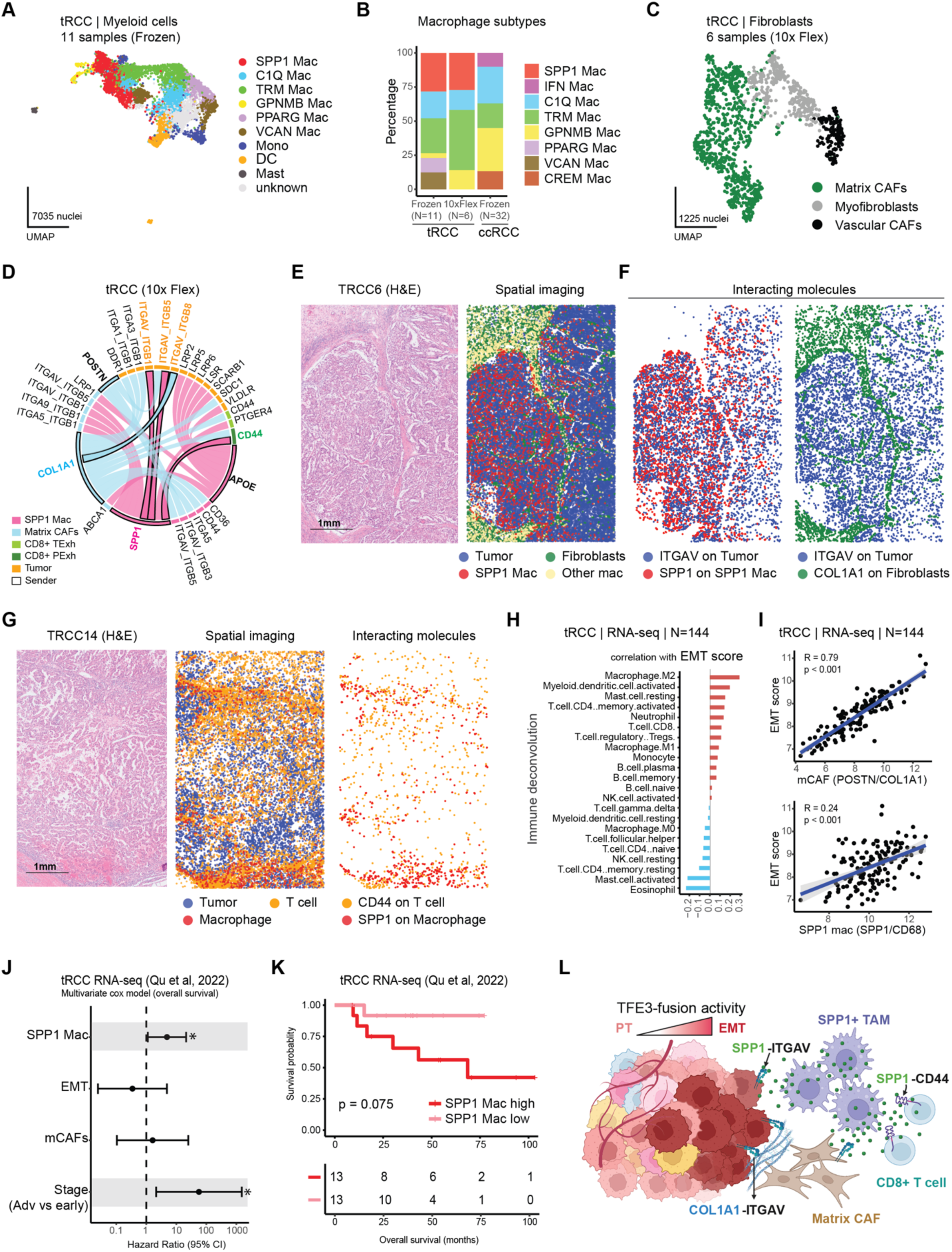
A suppressive tumor microenvironment in tRCC. A. UMAP depicting myeloid cell subtypes identified in the tRCC tumors from the frozen cohort. B. Stacked bar plot of macrophage subtypes in tRCC vs. ccRCC. C. UMAP of fibroblast subtypes identified in the tRCC (10x flex) cohort. D. Chord diagram showing inferred interactions of SPP1 macrophages with Matrix CAFs, tumor cells and CD8 T cell subtypes in the tRCC (10x flex) cohort. E. Hematoxylin and eosin (H&E) staining (left) and spatial transcriptomics (Nanostring CosMX, right) of a section of tRCC tumor from sample TRCC6. Tumor cells, fibroblasts, SPP1+ macrophages and other macrophages are highlighted in the spatial image plot. F. Spatial plots highlighting distribution of interacting molecules: SPP1 on SPP1+ macrophages with ITGAV on tumor cells (left) and COL1A1 on fibroblasts with ITGAV on tumor cells (right). G. H&E staining (left) and spatial imaging (Nanostring CosMX, middle) of a section of tRCC tumor from sample TRCC14. Tumor cells, T cells and macrophages are highlighted in the spatial image plot. Spatial plot highlighting distribution of interacting molecules: SPP1 on macrophages and CD44 on T cells (right). H. Correlation between sample-wise immune cell deconvolution scores from tRCC bulk RNA-seq data with corresponding EMT scores. I. Scatterplots showing correlation of EMT score with Matrix CAFs (top) and SPP1 Macrophages (bottom) in tRCC bulk RNA-seq cohorts. J. Multivariate cox model on overall survival from tRCC samples and association with SPP1 macrophages, EMT, matrix CAFs, and tumor stage (advanced vs early) as predictors. Significant predictors are highlighted. * indicates p-value <= 0.05. K. Kaplan-Meier survival curve for SPP1 low and SPP1 high tRCC samples in Qu et al. dataset. P-value was calculated by a pairwise log-rank test. L. Schematic illustrating suppressive tumor microenvironment (TME) in tRCC. Tumor cells exist in distinct transcriptional states, with TFE3-fusion driving repression of the epithelial identity and promoting transition to a mesenchymal state. Key suppressive interactions highlighted in this study are highlighted, including SPP1–ITGAV signaling from macrophages to tumor cells, SPP1–CD44 signaling from macrophages to CD8+ T cells, and COL1A1–ITGAV signaling from matrix CAFs to tumor cells.

Recent scRNA-seq studies have facilitated the identification of myriad cancer-associated fibroblast (CAF) subpopulations, which differ in transcriptional state and function^101,102^. We observed that tRCCs exhibited multiple fibroblast types including vascular CAFs, matrix CAFs (mCAFs), and myofibroblasts (**Figure 5C & S5D**). Notably, the two latter fibroblast types have also both been linked to a suppressive TME: mCAFs have been linked to the recruitment and differentiation of SPP1+ macrophages^103^ while myofibroblasts have been linked both to M2-polarization of macrophages and promotion of a desmoplastic stroma^104–106^.

Together, these findings point to a suppressive tumor microenvironment in tRCC, which may constrain response to immunotherapy despite adequate tumor-associated antigens, in part through the accumulation of SPP1+ macrophages that contribute to T cell dysfunction via impaired antigen presentation, and secretion of osteopontin that reinforces a non-permissive stromal rich immune niche^98,107^.

### A crosstalk of suppressive signals constrains immune-responsiveness in tRCC

To better define the crosstalk between malignant, immune, and stromal compartments in tRCC, we performed a ligand-receptor interaction analysis across our tRCC snRNA-seq cohorts. This revealed a dense network of immunosuppressive interactions, with SPP1+ macrophages and mCAFs acting as central hubs (**Figure 5D & S5E; Methods**). Prominent candidate interactions included SPP1 on macrophages to ITGAV on tumor cells, COL1A1 on mCAFs to ITGAV on tumor cells, POSTN on mCAFs to ITGB3 on macrophages, and SPP1 on macrophages to CD44 on progenitor-exhausted CD8+ TILs. All of these putative interactions have been linked to the suppression of anti-tumor T-cell responses or to limiting the efficacy of immune therapies^107–117^ (**Figure 5D & S5E**).

To assess how these interactions were spatially organized within the tumor architecture, we performed single-cell spatial transcriptomics using the Nanostring CosMx platform on two tRCC specimens (TRCC6 and TRCC14, **Figure S5F-H**). Spatial ligand-receptor mapping confirmed the presence of SPP1-ITGAV and COL1A1-ITGAV interactions at points of macrophage-tumor and fibroblast-tumor interaction, respectively, in the TRCC6 sample (**Figure 5E-F**). We also observed that EMT-high tumor cells were in proximity to SPP1+ macrophages in this sample (**Figure S5G**). In TRCC14 (the sample with highest T-cell infiltration on snRNA-seq, **Figure S4E**), we additionally identified spatial proximity between SPP1+ macrophages and CD44+ CD8⁺ T cells, a known suppressive axis implicated in T cell exhaustion^115^ (**Figure 5G**). Moreover, these associations were validated in a bulk RNA-seq dataset of 95 tRCC tumors, where an EMT signature score positively correlated with signatures for M2-like macrophages (by immune deconvolution), mCAFs, and SPP1+ macrophages (**Figure 5H-I**).

Finally, to evaluate the clinical consequences of our findings, we determined the association between overall survival and RNA signatures of EMT, SPP1+ macrophages (SPP1/CD68) and mCAFs (COL1A1/POSTN) in a tRCC bulk RNA-seq cohort^23^. Via Cox multivariable regression incorporating tumor stage, SPP1+ macrophage score and advanced stage emerged as independent predictors of decreased survival in tRCC (**Figure 5J-K**).

Overall, these results support the notion that an EMT program in tRCC malignant cells contributes to an immunosuppressive microenvironment through spatially structured interactions between EMT-activated tumor cells, SPP1+ macrophages, mCAFs, and CD8+ TILs (**Figure 5L**).

## DISCUSSION

By integrating snRNA-seq with snATAC-seq, spatial transcriptomics, and TCR-seq on a cohort of 20 tRCC samples from 15 patients, we present a detailed molecular view of tRCC. Our study identifies conserved malignant transcriptional programs in tRCC, clarifies the cellular origin of this cancer, and elucidates how activity of the TFE3 driver fusion remodels the immune microenvironment. Additionally, we validate our key results against 128 tRCC bulk transcriptomes and perform a detailed comparative analysis with ccRCC (the most well-studied kidney cancer), revealing significant molecular differences. To our knowledge, this represents the most exhaustive single-cell study performed to date on this rare cancer and offers a framework for mechanism-inspired target discovery in this aggressive RCC subtype.

We observed six conserved MPs in tRCC, which mostly overlap with oncogenic MPs identified in ccRCC or other cancers, with some key distinctions. First, while PT and EMT MPs are identified in both tRCC and ccRCC, their utilization differs substantially; the EMT MP is more prevalent in tRCC malignant cells, whereas the PT program is more dominant in ccRCC. The induction of a mesenchymal phenotype in tRCC appears to be directly linked to TFE3-fusion activity (**Figure 3C-D**)^23,28^. Additionally, some tRCC malignant cells express an angiogenesis MP, notably in the absence of *VHL* inactivation-associated HIF2a signaling observed in ccRCC^3–6^. This suggests alternative mechanisms of neovascularization in tRCC, possibly mediated by inflammatory cytokines, as evidenced by co-enrichment of TNF and inflammatory MPs with the angiogenesis MP^61,62^ (**Figure 3D**). This observation may also explain the higher response rates to immune checkpoint inhibitor (ICI)-VEGF inhibitor combination therapies compared to ICI alone^17^.

Despite the genomic homogeneity of tRCC, which is characterized by a low mutational burden and relatively few recurrent alterations beyond the driver fusion^12,21–23^, we observed significant transcriptional heterogeneity both within and among tumors. Similar non-genetic heterogeneity has been described in other fusion-driven cancers, including Ewing sarcoma^118,119^, alveolar rhabdomyosarcoma^120,121^, and synovial sarcoma^122^, where variation in fusion oncogene activity gives rise to distinct cellular states associated with metabolism, differentiation, or immune evasion. This transcriptional plasticity has also been identified for c-Myc – which, like TFE3, is an oncogenic transcription factor in the bHLH-LZ family^123^. In tRCC, cell-to-cell variability in TFE3 fusion activity may stem from heterogeneity in the expression levels of the fusion itself, variability in the expression of key co-factors or differences in chromatin accessibility.

Our single-cell analyses indicate that despite different genetic drivers and transcriptional programs, tRCC and ccRCC both originate from VCAM1-expressing proximal tubule cells. Interestingly, other origins for tRCC, such as parietal epithelial cells, were predicted in a few cases. While these instances could represent imprecise cell type classification, an alternative explanation is that tRCC has the capacity to arise from multiple potential cell types that are permissive to generate the driver fusion. The presence of *TFE3* fusions in non-kidney cancers (e.g., ASPS, PEComa) further supports their potential to emerge from diverse cells^124–129^. Additionally, it is plausible that the histological plasticity observed in tRCC - which can microscopically resemble both clear-cell and papillary subtype RCCs - may be rooted in its ability to arise in diverse cells of origin.

Our study provides mechanistic insight into the clinical observation of suboptimal responses to ICI in tRCC^12,17,76,130^. The TME in tRCC contains immunosuppressive cells like SPP1+ TAMs and COL1A1+ mCAFs, which collaborate to suppress the anti-tumor activity of CD8+ TILs^101,102^. In addition to a suppressive SPP1+ TAM/CD8+ T-cell interaction^115^, SPP1 secreted by TAMs activates fibroblasts, amplifying matrix deposition^107,116,117,131–134^ and further immune suppression^96–98,135^. Notably, these suppressive populations have been linked to a tumor-intrinsic EMT program in other cancers^136–141^, which we show is also activated in tRCC downstream of the driver fusion. Thus, despite potential neoantigens derived from the TFE3 fusions, a suppressive TME may constrain anti-tumor immunity in tRCC. Interestingly, however, ASPS – which also harbors a TFE3 driver fusion – was shown to have a 37% objective response rate to the single-agent PD-L1 inhibitor, atezolizumab, in a recent clinical study^142^. A comparative analysis of tRCC and ASPS may offer an opportunity to disambiguate microenvironmental factors from oncogenic drivers alone in determining immune resistance. Moreover, targeting the EMT-macrophage-fibroblast circuit may offer strategies to reprogram the TME and potentiate immune therapies in tRCC.

## ACKNOLWEDGEMENTS

S.R.V. acknkowledges the Doris Duke Charitable Foundation (Clinician-Scientist Development Award grant no. 2020101), Department of Defense Kidney Cancer Research Program (DoD KCRP) (W81XWH-22-1-016), Damon Runyon-Rachleff Innovation Award (grant no. 71-22), the National Cancer Institute (NCI) R01CA279044, NCI R01CA286652, V Scholar Foundation (V2022-018) and Rally Foundation (23IN37). T.K.C. is supported in part by the Dana-Farber/Harvard Cancer Center Kidney SPORE (2P50CA101942-16) and Program 5P30CA006516-56, the Kohlberg Chair at Harvard Medical School and the Trust Family, Michael Brigham, Pan Mass Challenge, Hinda and Arthur Marcus Fund and Loker Pinard Funds for Kidney Cancer Research at DFCI. S.S. was supported in part by the NCI Dana-Farber/Harvard Cancer Center Kidney Cancer SPORE (P50-CA101942-12) and the NCI Dana-Farber/Harvard Cancer Center Kidney Program (P30 29 CA06516). E.M.V.A. acknowledges funding from NIH R01CA278980. C.-Z.Z. received funding from Claudia Adams Barr Program for Innovative Cancer Research, Breakthrough Cancer, NCI (K22CA216319 and R01CA286652). D.J.E reports funding from P50 CA254861-01A1 SPORE in Prostate Cancer. P.K. received funding from DoD KCRP Postdoctoral and Clinical Fellowship (HT94252310066). C.N.W. received funding from AACR-Exelixis Renal Cell Carcinoma Research Fellowship (24-40-66-WEIS). J.L. received funding from DoD KCRP Postdoctoral and Clinical Fellowship (W81XWH-22-1-0399).

We are deeply grateful to the patients and their families for their generosity in contributing to this research project. Single-nucleus RNA/ATAC sequencing were performed at the Center for Cancer Genomics at Dana-Farber Cancer Institute. We acknowledge contributions from Mark Connelly, Nina Listopadzki, and Julius Nevin with the single-nucleus RNA/ATAC sequencing. TCR sequencing was performed at the Translational Immunogenomics Lab at Dana-Farber Cancer Institute. We acknowledge contributions from Kenneth Livak, Shuqiang Li, and Haoxing Lyu with the TCR sequencing. Spatial imaging was performed at the Center for Cellular Profiling at Brigham Women’s Hospital. We acknowledge contributions from Ce Gao, Junning Case, and Miles Tran with the spatial imaging. Graphical illustrations in Figures 1A, 5L, S2C, S3B were created with BioRender.com.

## AUTHOR CONTRIBUTIONS

Conceptualization: P.K. and S.R.V. Performed or designed bioinformatic analyses: P.K. and S.R.V with assistance from C.N.W., Y.C., J.W., R.D., J.L., S.Y.C., C-Z.Z., E.M.V.A. Procurement, processing, and/or histopathologic analyses of clinical samples: T.K.C., C.N.W., J.L.H., D.J.E., E.M.V.A, J.H., S.S., S.M., Y.L.N., A.B.S. Processing of samples for single-nucleus RNA/ATAC: A.R.T., A.N., and Y.R. Supervision and funding acquisition: S.R.V. and T.K.C. Manuscript, original draft: P.K. and S.R.V. All authors edited and approved the final draft of the manuscript.

## METHODS

### Study approval and sample collection

A total of 20 tRCC tumors from 15 individuals were collected and sequenced using one or more of the following platforms: 10x Genomics 3’ snRNA-seq, 10x 3’ multiome (RNA + ATAC), 10x-Flex fixed RNA-seq, bulk TCR sequencing, and spatial transcriptomics. For subject TRCC18, the primary tumor was collected during initial nephrectomy, and 13 metastatic sites were collected during rapid autopsy as previously described, 3 of which were subjected to snRNA-seq^21^ (see **Figure S3**). The cohort included 8 females and 7 males. Sample preservation methods (fresh frozen or FFPE-fixed) and additional clinical details are provided in **Table S1** & **Figure S1A**. All individuals provided written informed consent for the use of their samples in sequencing and research. The rapid autopsy for subject TRCC18 was performed under a genitourinary oncology rapid autopsy protocol approved by the Dana-Farber/Harvard Cancer Center Institutional Review Board (IRB).

### Single-nucleus RNA sequencing (3’ RNA and 3’ multiome)

From fresh frozen tissues, nuclei isolation was performed as previously described^143^, using low-retention microcentrifuge tubes (Fisher Scientific, Hampton, NH, USA) throughout the procedure to minimize nuclei loss. Briefly, OCT-embedded tissue fragments were rapidly thawed on wet ice to remove excess OCT, mechanically dissociated by pipetting, and homogenized in TST solution. The homogenate was filtered through a 30 µm MACS SmartStrainer (Miltenyi Biotec, Germany) and pelleted by centrifugation for 4 minutes at 500 × g at 4°C. The resulting nuclei pellet was resuspended in 150 µL of 10x Genomics Diluted Nuclei Buffer. Trypan blue-stained nuclei were counted manually using INCYTO C-Chip Neubauer Improved Disposable Hemacytometers (VWR International Ltd., Radnor, PA, USA).

For 3’ RNA-seq, approximately 8,000 nuclei per sample were loaded per channel of a Chromium Next GEM Chip G and processed on the Chromium Controller (10x Genomics, Pleasanton, CA, USA), followed by cDNA synthesis and library construction according to the manufacturer’s protocol (Chromium Next GEM Single Cell 3ʹ Reagent Kits v3.1 User Guide, Rev D). Libraries were normalized and pooled for sequencing on a NovaSeq SP-100 flow cell (Illumina, Inc., San Diego, CA, USA).

For multiome (RNA + ATAC) profiling, approximately 16,000–25,000 nuclei per sample were loaded per channel of a Chromium Next GEM Chip J and processed similarly using the Chromium Controller, followed by simultaneous transposition, cDNA synthesis, and library preparation as per the manufacturer’s instructions (Chromium Next GEM Single Cell Multiome ATAC + Gene Expression User Guide, Rev F). Libraries were normalized and pooled for sequencing across two NovaSeq SP-100 flow cells (Illumina, Inc.).

### Single-nucleus RNA sequencing (10x flex fixed)

FFPE tissue processing and RNA isolation were performed using the RNeasy FFPE Kit (Qiagen) on two 10-μm scrolls per sample, following the manufacturer’s protocol. RNA quality was assessed via DV200 analysis using the Agilent 2100 Bioanalyzer. For nuclei isolation, three 30-μm scrolls per sample were deparaffinized in xylene, rehydrated through graded ethanol washes, and incubated in Milli-Q water followed by cold PBS. Tissue dissociation was performed using enzymes from the Miltenyi FFPE Tissue Dissociation Kit and Liberase™ TM, followed by mechanical dissociation with the gentleMACS Octo Dissociator. Nuclei were filtered, centrifuged, and resuspended in 10x Genomics quenching buffer, and counted using AO staining with a Cellometer-K2 Fluorescent Cell Counter. All steps were performed independently for each of the nine samples.

Approximately one million nuclei per sample were hybridized with Human WTA Probes (10x Genomics) for 23 hours at 42°C, washed three times per the manufacturer’s protocol (CG000527-Rev E), and counted. Around 50,000 nuclei per sample were pooled, filtered, and assessed for quality. A total of ∼144,000 pooled nuclei were processed using the Chromium Next GEM Single Cell Fixed RNA for Multiplexed Samples workflow. GEM formation, reverse transcription, cleanup, and pre-amplification were performed according to the user guide, followed by FLEX-GEX library construction. Final libraries were quantified on the Agilent Bioanalyzer 2100.

### Single-nucleus RNA sequencing data processing and analysis

Sequencing data were demultiplexed, aligned, and quantified using the 10x Genomics Cell Ranger pipeline (v3.1.0). Reads were aligned to the GRCh38 reference genome (Gencode annotation), and gene-by-cell UMI count matrices were generated. Count matrices were loaded into R using the Read10X function and processed using the Seurat v4^144^ package. Initial quality control retained features expressed in at least 3 cells, and cells with >200 and <3,000 detected features. Cells with >5% mitochondrial gene content and those flagged as potential doublets using the Scrublet^145^ Python package (default parameters) were excluded.

Filtered data were normalized using Seurat’s NormalizeData function (scaling factor = 10,000), and the top 2,000 variable features were identified per sample using FindVariableFeatures. All genes were used for scaling (ScaleData) and principal component analysis (PCA). PCs were manually reviewed per sample based on variance explained and top feature loadings to guide downstream dimensionality reduction and clustering. Clustering was performed using the Louvain algorithm at multiple resolutions (0.2–1.2); resolution was selected manually based on marker gene expression patterns within subclusters.

Broad lineage identities (tumor, lymphoid, myeloid, endothelial, fibroblast, normal epithelial) were assigned to clusters using canonical marker genes (e.g., Tumor: *TRIM63*, *GPNMB*, *NMRK2*, *CTSK*; Myeloid: *SLC8A1*, *CD74*, *CD163*, *CTSB*; Lymphoid: *THEMIS*, *PTPRC*, *IKZF1*, *CD96*; Endothelial: *PECAM1*, *FLT1*, *VWF*, *PTPRB*; Fibroblasts: *COL3A1*, *COL1A1*, *FBN1*, *LUM*). An additional reference-based cell annotation was performed using SingleR^146^ with the Blueprint-ENCODE reference. For the normal kidney sample (TRCC19N), nephron-specific cell types were annotated using marker genes from Muto et al^53^.

For merged analysis across all samples, individual Seurat objects were combined using MergeData without batch correction. The merged dataset was reprocessed to identify highly variable genes, followed by PCA, UMAP, and clustering. For tumor-specific subcluster integration (Figure 3), batch correction was performed using Harmony^147^. Module scores were computed using the AddModuleScore function across all snRNA-seq analyses.

Differentially expressed genes between clusters were identified using FindAllMarkers with the following criteria: expression in ≥10% of cells within a cluster and a log_2_ fold-change >0.25, yielding a total of 2,161 marker genes (**Figure 1C**). To assess tumor cell-of-origin (**Figure S2**), FindTransferAnchors and TransferData functions were used to map tumor cells to nephron epithelial subtypes. Cell–cell communication analysis (**Figure 5 & S5**) was performed using the LIANA^148^ framework with default parameters, incorporating multiple tools (NATMI, Connectome, logFC, SCA, CellPhoneDB), and prioritizing ligand–receptor interactions with a consensus aggregate rank ≤ 0.05.

### Tumor Meta-Program NMF analysis

Non-negative matrix factorization (NMF) analysis was performed separately for malignant cells from each tumor sample. For each tumor, raw gene expression counts were normalized using Seurat’s NormalizeData() function with default parameters. Highly variable genes (HVGs) were identified using FindVariableFeatures() (top 2,000 genes), and their expression values were center-scaled using ScaleData(), regressing out the percentage of mitochondrial gene expression. Any negative values in the scaled expression matrix were set to zero.

NMF was then applied to the scaled HVG expression matrix using the ‘nsNMF’ method (implemented in the NMF R package v0.23.0), across a range of factorization ranks from 5 to 25, as described previously^63^. To define robust gene modules, we implemented a previously described ranking strategy^41^: first, we generated two ranking matrices: (1) for each factor, genes were ranked by contribution weight; (2) for each gene, contributing factors were ranked by weight. A gene was included in a module if it ranked highest for a given factor, and genes were added sequentially until one was encountered that ranked higher for another factor. Modules with fewer than five genes were discarded, and this process was iteratively applied for each NMF rank.

To select an optimal rank, we identified the highest rank at which the number of resulting gene modules equaled the rank, ensuring stability and interpretability. The resulting gene modules across individual tumors were then filtered to retain only those with at least 5% overlap (Jaccard index) with two or more modules from other tumors, indicating shared transcriptional programs. Modules with fewer than five genes or those lacking significant enrichment for known gene sets were excluded from further analysis.

For biological annotation of the remaining gene modules, we performed pathway and ontology enrichment analysis using the ClusterProfiler^149^ and EnrichR^150^ packages in R. Enrichment was assessed against Gene Ontology (GO) terms, KEGG pathways, Reactome pathways, Hallmark gene sets (MSigDB), and PanglaoDB cell type markers.

To assess the distribution of meta-programs across individual samples, each malignant cell was annotated with the NMF module for which it exhibited the highest module score, calculated using Seurat’s AddModuleScore() function. To facilitate cross-cohort comparisons, tumor subclusters within the tRCC dataset were annotated with corresponding meta-programs that showed dominant activity within each subcluster. These annotations were based on enrichment of module scores and expression coherence. To compare our NMF-derived programs with previously published transcriptional programs, we retrieved gene signatures from published supplementary datasets and calculated module scores for these signatures across our NMF clusters.

### snRNA-seq Regulon analysis

To infer transcription factor (TF) regulons and quantify their activity across tumor NMF clusters from snRNA-seq data, we applied the pySCENIC^151^ pipeline (v0.12.1) on subsetted tumor cells from merged object. Gene co-expression modules were first generated using the grn step, based on the filtered expression matrix and a curated hg38-based list of human TFs. These initial co-expression modules were then refined during the ctx step by evaluating motif enrichment in regulatory regions located +/-10 kb from transcription start sites, using the hg38 motif ranking database.

To ensure biological relevance, only regulons with at least five target genes, a normalized enrichment score (NES) ≥ 2, and an area under the curve (AUC) threshold ≥ 0.1 were retained. Dropout masking was applied during this step to mitigate the effects of sparsity in single-nucleus RNA-seq data. Subsequently, regulon activity scores were calculated on a per-cell basis using the aucell step, producing a cell-by-regulon AUC matrix.

Differential regulon activity between epithelial-to-mesenchymal transition (EMT)-like and proximal tubule (PT)-like tumor subpopulations, defined based on tumor subclusters in Figure 3D, was assessed using the FindMarkers function in Seurat. Regulons were ranked based on their average log fold change and adjusted p-value within EMT and PT clusters to identify TFs potentially driving lineage-specific transcriptional programs.

### snATAC-seq data analysis

All snATAC-seq samples were initially processed using Cell Ranger ATAC (v2.0, 10x Genomics) with default parameters, aligning to the GRCh38 reference genome (refdata-cellranger-arc-GRCh38-2020-A-2.0.0). Output barcode and fragment files were subsequently analyzed using the Signac^152^ R package (v1.12.0). Peak calling was performed with the CallPeaks() function using MACS2^153^. Quality control thresholds were applied based on the following parameters: nCount_RNA (1000 < count < 25000), blacklist_fraction, nucleosome_signal, and TSS.enrichment, each filtered at +/-3 median absolute deviations from the median per sample. Following QC, peak counts were normalized using TF-IDF normalization and dimensionality reduction was performed via Singular Value Decomposition (SVD) using latent semantic indexing (LSI, components 2–30). Non-linear dimensionality reduction and clustering were then performed on the LSI components. Cluster annotation was carried out based on known chromatin accessibility markers and label transfer from matched snRNA-seq data. For integrated analyses, samples were harmonized using Harmony to correct for batch effects.

Differential chromatin accessibility between tRCC and ccRCC tumor cells was assessed using the FindMarkers() function in Signac. Genomic annotation of differentially accessible peaks was performed using the ClosestFeature() function to associate peaks with the nearest genes. Logistic regression was implemented to account for differences in sequencing depth across samples.

Motif enrichment analysis was then performed using AddMotifs() and the JASPAR2020 motif database. Top differentially accessible peaks were selected based on thresholds of |avg_log2FC| ≥ 1 and adjusted p-value ≤ 0.05. Overrepresented motifs in tRCC and ccRCC tumor cells were identified using the FindMotifs() function.

### CosMx Spatial data analysis

Five-micron (5 µm) FFPE tissue sections were mounted on Leica BOND PLUS slides (Leica Biosystems, Cat. S21.2113.A). Sample preparation followed the manufacturer’s protocol for the Human Universal Cell Characterization 1000 Plex Panel (Nanostring, Part #122000157) as outlined in the CosMx SMI Manual Slide Preparation Manual (MAN-10159-01). Slides were initially baked overnight at 60°C to enhance tissue adherence. Subsequent sample processing steps included sequential deparaffinization, target retrieval (15 min at 100°C), and permeabilization with proteinase K (3 µg/mL for 15 min at 40°C). Fiducials were then applied, followed by post-fixation and NHS-acetate treatment. Tissue sections were hybridized with denatured probes from the universal panel and the default add-on panel for 18 hours at 37°C.

After hybridization, slides were washed and stained with DAPI (15 min at room temperature) and a marker stain mix containing PanCK, CD45, CD68, and CD298/B2M (for cell segmentation). Slides were washed again and loaded onto the CosMx SMI instrument (Machine ID: CosMx_0020; Serial Number: INS2301H0020) for UV bleaching, cyclic imaging, and scanning. Raw images were processed and decoded using the default pipeline on Atomx SIP, a cloud-based service. The resulting Nanostring spatial transcriptomics data were imported into R and analyzed using the Seurat package. Cell type annotation was performed by label transfer from matched single-nucleus RNA-seq datasets and further validated by differential gene expression analysis using the FindMarkers() function. EMT program was identified based on module score analysis and cells with positive scores were considered to be EMT positive.

### Bulk-RNA data analysis

Data from external tRCC cohorts were obtained as previously described^12^ for TCGA^32^, Motzer et al.^33^, Wang et al^34^. Raw tRCC sequencing data and metadata for Sun et al^22^ and Qu et al^23^ were obtained from the respective studies. Reads were aligned to the human reference genome (GRCh38.p13 assembly) using STAR (v2.7.10a) and quantified as paired-end reads against the GENCODE v45 transcript reference using RSEM (v1.3.1). Expression matrices from each dataset were log-transformed, centralized, and combined to reduce batch-related differences prior to downstream analysis. For immune deconvolution analysis, we used TIMER2.0, and the results plotted are from CIBERSORTX. Expression scores for different meta-programs were identified by calculating average expression of genes comprising each meta-program (**Table S3**).

To evaluate the clinical relevance of transcriptional programs and immune cells, we used gene expression and survival data from Qu et al^23^. to calculate program scores and assess their association with patient outcomes (**Figure 5**). Program scores were derived as the average expression of genes comprising each meta-program after log-transformation and centralization of the expression matrix. We performed multivariate survival analysis using Cox proportional hazards models (coxph function, survival R package), adjusting for clinical and molecular covariates including SPP1 macrophage signature (*SPP1*/*CD68*), EMT status (based on NMF signature; **Table S3**), matrix CAFs (*COL1A1*/*POSTN*), and tumor stage (categorized as early [I-II] vs. advanced [III-IV]). Kaplan-Meier curves were generated to compare overall survival (OS) between stratified groups, SPP1 macrophage high vs. low (dichotomized at the median), and statistical significance was evaluated using the log-rank test (survminer v0.4.9).

### tRCC vs ccRCC comparative data analysis

For comparative transcriptomic analysis between tRCC and ccRCC, we utilized both single-nucleus RNA sequencing (snRNA-seq) and bulk RNA-seq datasets. For the snRNA-seq analysis, we subsetted tumor cells from tRCC and ccRCC samples; where ccRCC data were obtained from Wu et al.^35^, processed in alignment with their published pipeline, and restricted to the cells included in their original study. The resulting Seurat objects (from fresh frozen samples) were merged, and differential gene expression analysis was performed using the FindMarkers function in Seurat, applying the Wilcoxon rank-sum test. Genes with an average log fold change ≥ 1.5 and adjusted p-value < 0.05 were considered significantly differentially expressed.

For bulk RNA-seq data, expression matrices from each dataset^22,32,33^ were log-transformed and centralized. These normalized datasets were then combined, and differential expression between tRCC and ccRCC was assessed by comparing gene-wise z-scores and Wilcoxon test with benjamini-hochberg correction. Genes with a z-score difference ≥ 0.5 and adjusted p-value < 0.05 were considered significant.

Pathway enrichment analysis was performed on the set of intersecting differentially expressed genes identified in both snRNA-seq and bulk RNA-seq platforms using ClusterProfiler^149^ (v4.13.3).

### TFE3-fusion transcriptional signature

The TFE3-fusion signature was derived using previously published bulk RNA-seq datasets from HEK-293T cells engineered to overexpress various TFE3 fusion constructs (ASPSCR1-TFE3, SFPQ-TFE3, NONO-TFE3, and PRCC-TFE3^12^) compared to mock-transfected controls, along with data from the tRCC cell line FUUR1^27^ following shRNA-mediated knockdown of the ASPSCR1-TFE3 fusion. Differentially expressed genes were first identified in the overexpression datasets by filtering for those with a fold change ≥ 2 and p-value ≤ 0.05 in at least two of the fusion constructs. This gene set was intersected with genes significantly downregulated upon TFE3 fusion knockdown in FUUR1 to define a consensus fusion-driven program, yielding a core list of 113 genes (**Table S3**). The resulting TFE3-fusion signature was then used to quantify TFE3 fusion activity across datasets: for snRNA-seq, scores were calculated per nucleus using the AddModuleScore function in Seurat; for bulk RNA-seq data, activity scores were computed as the average expression of the signature genes after log-normalization and centering.

### T cell receptor sequencing and analysis

T cell receptor (TCR) sequencing was performed using the rhTCRseq protocol for bulk TCR sequencing and repertoire analysis as previously described in Li et al^154^. For data analysis, raw FASTQ files were processed using an in-house rhTCRseq pipeline. Reads were first aligned to either the TRA or TRB locus using BLAST and subsequently assembled into TCR clonotypes using MiXCR. CDR3 sequences were extracted, and clonotypes were collapsed based on CDR3 similarity to generate a final list of unique clonotypes and their respective frequencies.

### Quantification and statistical analysis

Quantification and statistical analyses were performed as described in the corresponding figure legends and throughout the text. Where applicable, p-values, the statistical tests used to generate them, and the exact number of samples or nuclei (N/n) are indicated either in the figure legends or the main text.

---

**Figure S1.**
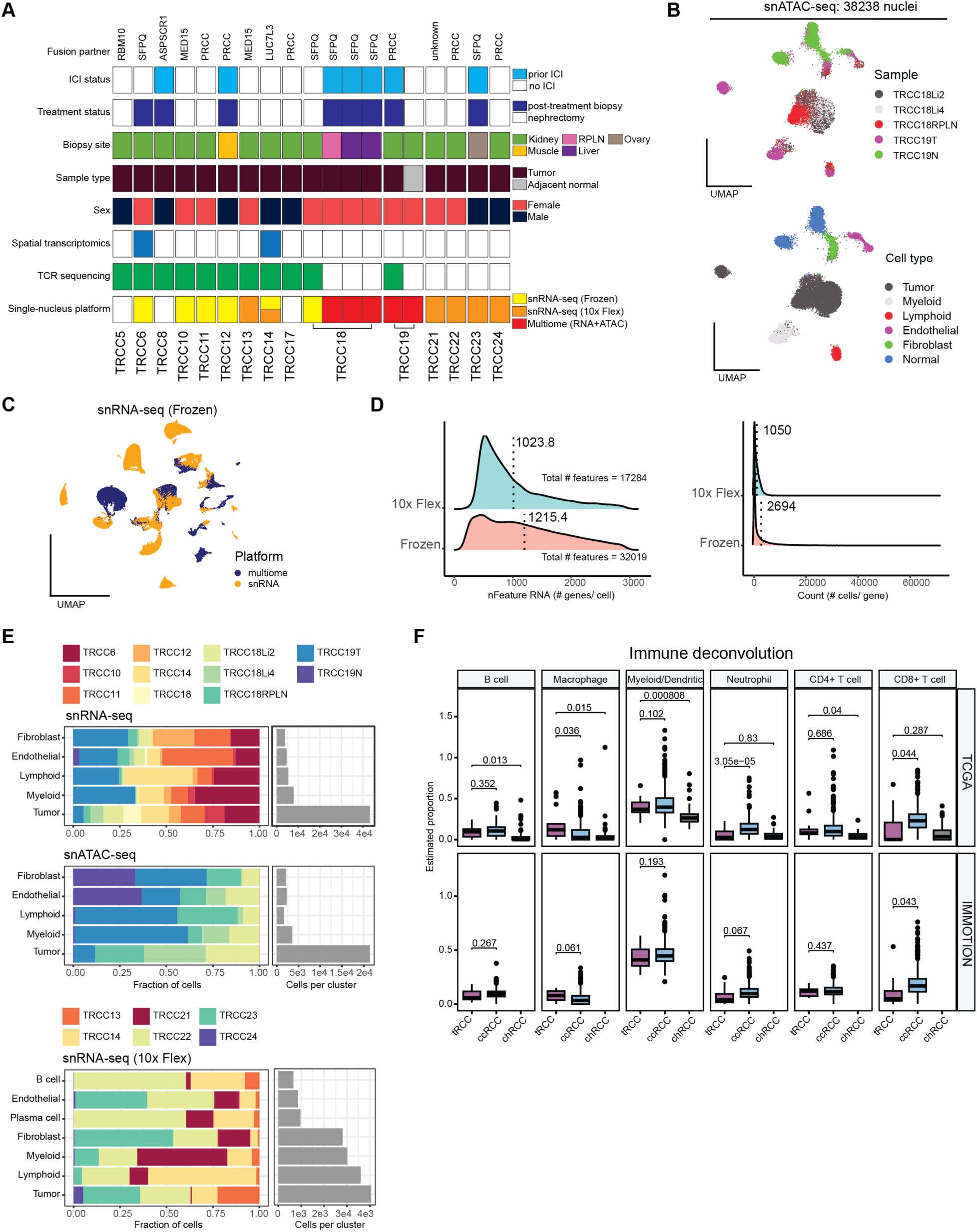
Characteristics of tRCC snRNA-seq cohorts. A. Summary of clinicopathological characteristics, treatment history, biopsy site, fusion partner, and molecular profiling approaches used for tRCC samples in this study. B. UMAPs of tRCC snATAC-seq data representing samples (top) and major cell lineages (bottom). C. UMAP representation of snRNA-seq data from the frozen cohort, highlighting samples processed via multiome (RNA+ATAC) or RNA-only platforms. D. Distribution of number of genes detected per nucleus in frozen and 10x flex fixed cohorts (left) and number of cells in which each gene was detected (right). E. Stacked bar plots showing lineage composition in three tRCC cohorts: snRNA-seq (Frozen), snATAC-seq, and snRNA-seq (10x Flex). Adjacent bar plots indicate total cell counts per lineage. F. Boxplots showing the proportions of immune cell types identified through bulk RNA-seq deconvolution in RCC subtypes along with p-values for indicated comparisons.

**Figure S2.**
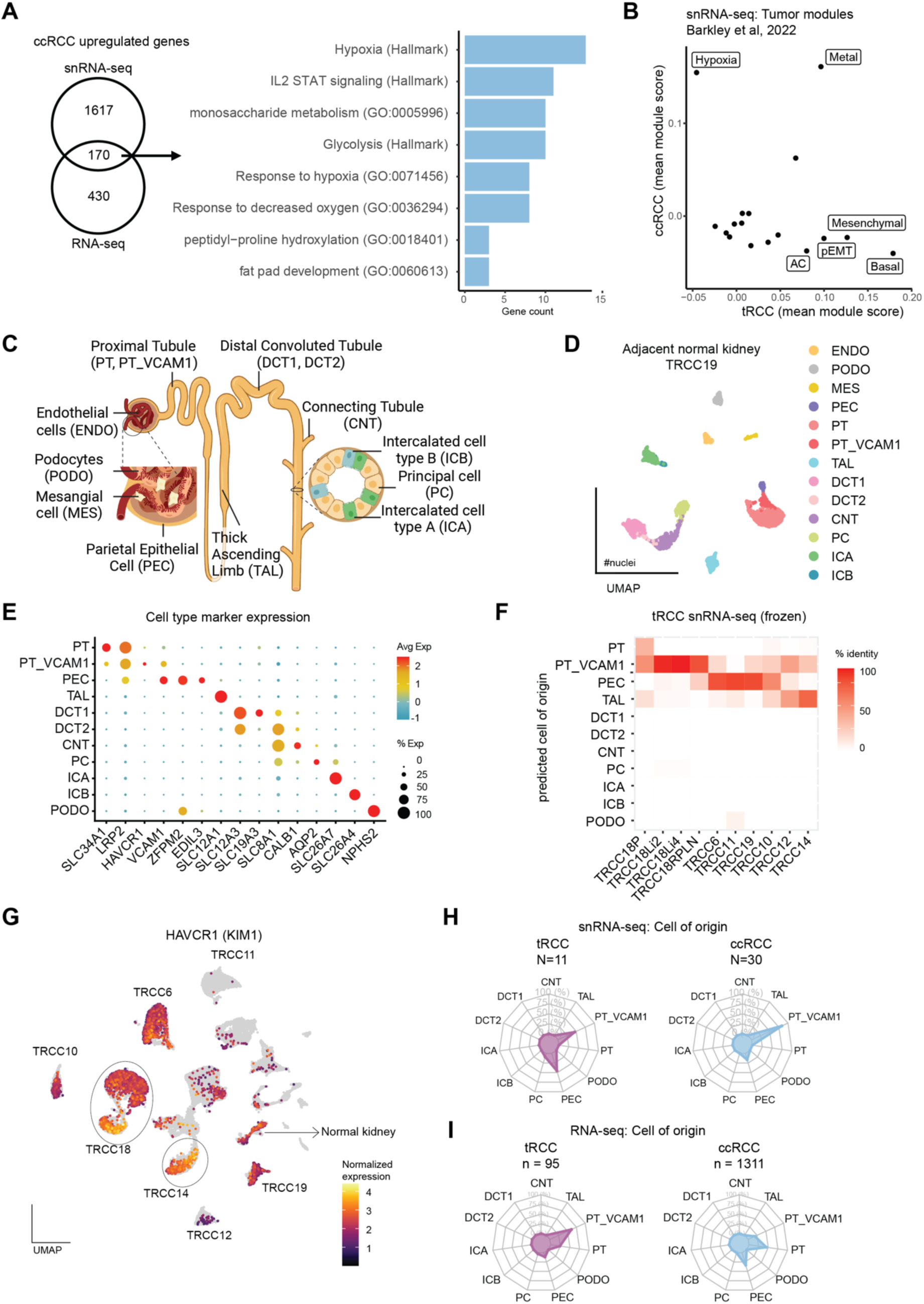
tRCC and ccRCC tumors arise from a similar cell of origin. A. Venn diagram depicting genes upregulated in ccRCCs vs tRCC in snRNA-seq and bulk RNA-Seq datasets (left). Bar plot with top enriched pathways in ccRCCs based on gene-ontology (right). B. Scatterplot of module scores in tRCC vs ccRCC tumor cells from snRNA-seq data, based on a published study^41^. C. Schematic of human nephron depicting different normal kidney cell types. D. UMAP from snRNA-seq of tumor-adjacent normal kidney sample, TRCC19N. E. Dot plot depicting the expression of marker genes in normal kidney cell types in TRCC19N. F. Heatmap depicting percent identity between tumor cells in individual tRCC samples and normal kidney cell types (tRCC snRNA-seq frozen cohort). G. UMAP showing expression of *HAVCR1* (KIM1) across tRCC tumor and non-tumor cells from the frozen cohort. H. Radar plots indicating the similarity of tRCC and ccRCC to different normal kidney cell types (snRNA-seq frozen cohorts). I. Radar plots indicating the similarity of samples from tRCC and ccRCC bulk RNA-seq cohorts to different normal kidney cell types.

**Figure S3.**
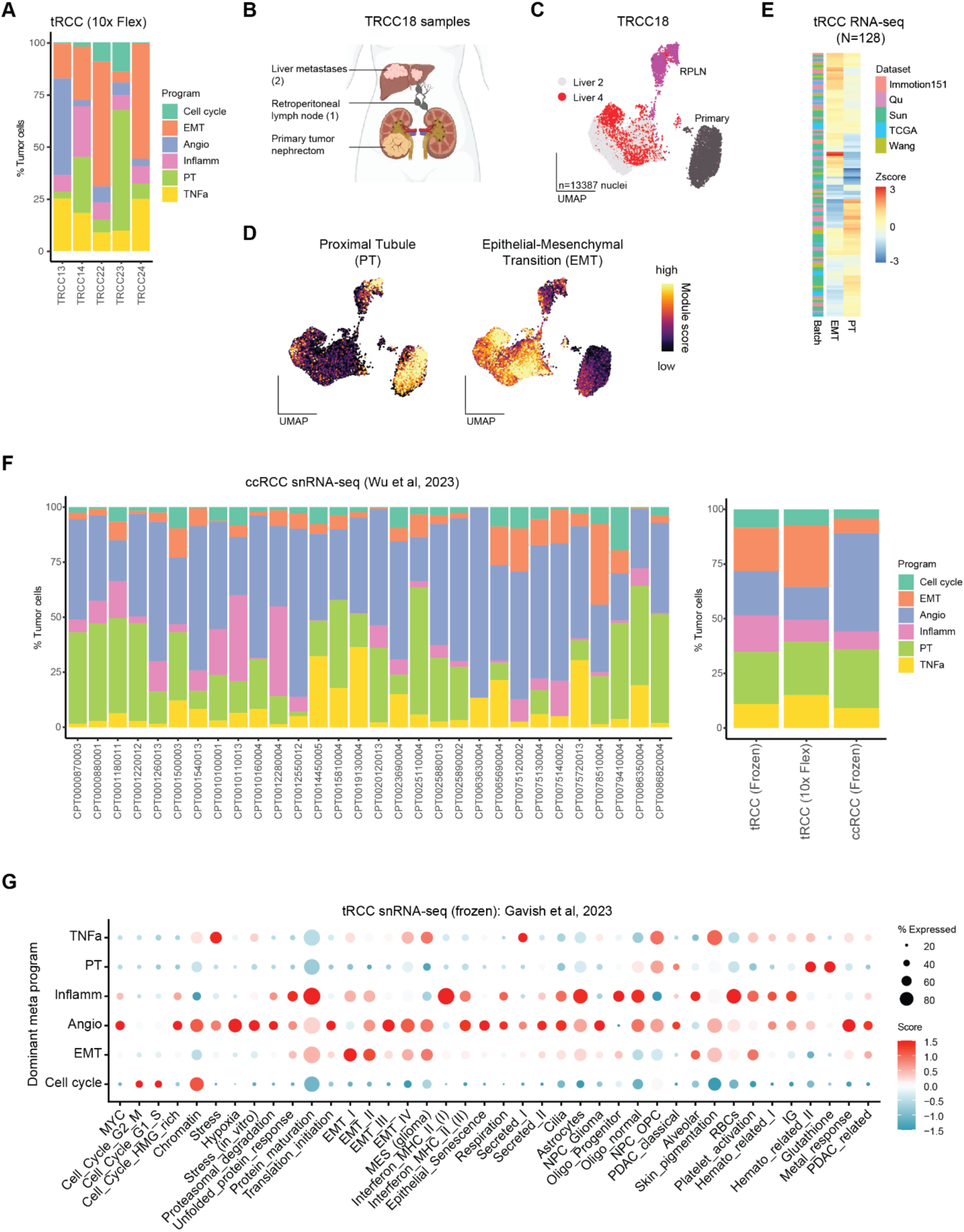
Characterization of tumor programs in tRCC and ccRCC. A. Stacked bar plot representing the distribution of tRCC NMF programs across the tRCC 10x Flex cohort. B. Schematic of samples related to primary and metastatic sites from patient TRCC18. C. UMAP depicting distribution of cells from TRCC18 by biopsy site. D. UMAP depicting proximal tubule (PT) and epithelial-mesenchymal transition (EMT) program scores across primary and metastatic sites of TRCC18. E. Heatmap depicting distribution of EMT and PT program scores across tRCC samples from the bulk RNA-seq cohort, annotated by dataset. F. Stacked bar plot representing the distribution of identified NMF programs across ccRCC snRNA-seq cohort (left) and comparison of overall proportions between all tRCC and ccRCC cohorts (right). G. Dot plot showing module scores of tumor NMF programs defined previously in a pan-cancer study by Gavish et al.^40^ and association with tRCC NMF programs identified in this study.

**Figure S4.**
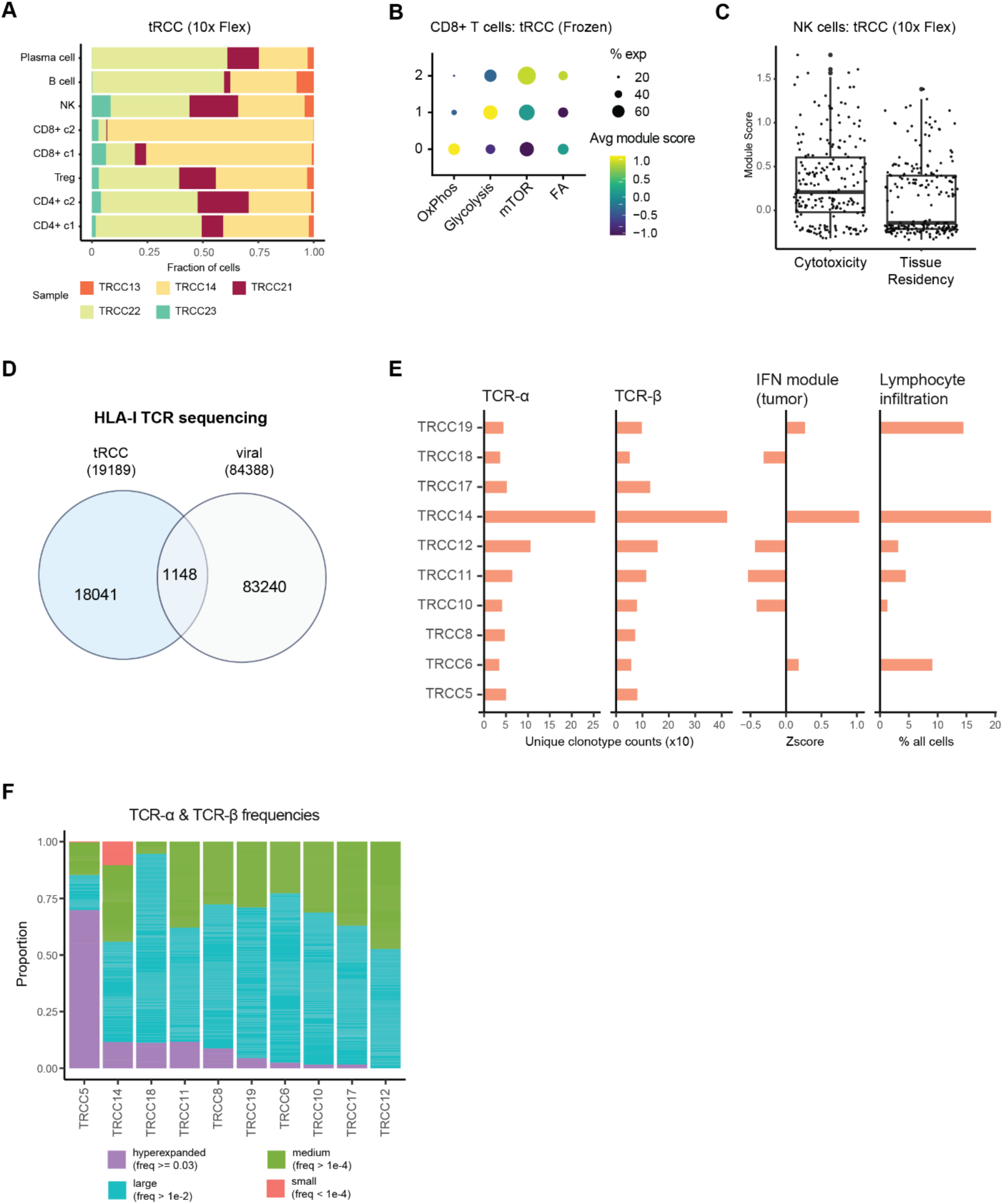
Lymphocyte infiltration and T cell receptor (TCR) clonality in tRCC. A. Stacked bar plot showing the composition of lymphocytes in tRCC 10x Flex cohort. B. Dot plot of metabolic pathway module scores in CD8 T cell subclusters from the tRCC frozen cohort. C. Box plot of cytotoxicity and tissue residency module scores in NK cells from the tRCC 10x Flex cohort. D. Venn plot showing intersection of TCR sequences from bulk-TCR⍺ and TCRβ sequencing of tRCC samples and publicly annotated viral TCR sequences. E. Bar plots of counts of unique TCR⍺ and TCRβ clonotypes identified in each tRCC sample, corresponding interferon (IFN) module scores in tumor cells, and percentage lymphocyte infiltration from snRNA-seq data. F. Proportions of TCR clonotypes in each tRCC sample based on their abundance: hyperexpanded, large, medium, and small. Proportions reflect values after removing overlapping viral sequences from D.

**Figure S5.**
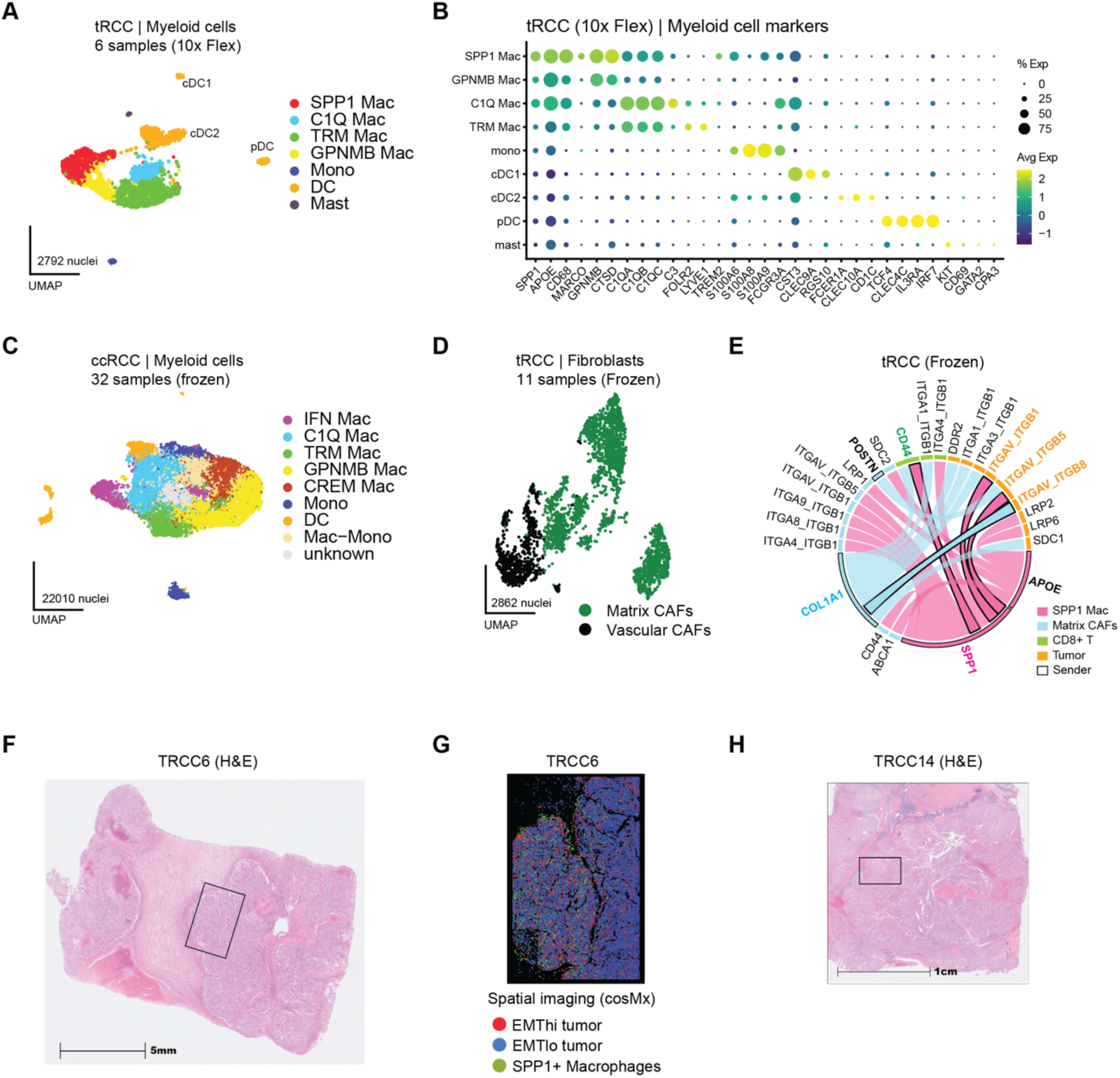
Suppressive myeloid cells and fibroblasts in the tRCC TME. A. UMAP of myeloid cell subtypes in the tRCC (10x Flex) cohort. B. Dot plot showing expression of marker genes in myeloid cell types in the tRCC 10x Flex cohort from A. C. UMAP depicting myeloid cell subtypes identified in the ccRCC (frozen) cohort. D. UMAP depicting fibroblast subtypes identified in the tRCC (frozen) cohort. E. Chord diagram showing interactions of SPP1 macrophages and Matrix CAFs (senders) with tumor cells and CD8 T cells in the tRCC (frozen) cohort. F. H&E staining of the full slide of sample TRCC6. The section highlighted indicates the field of view utilized for spatial imaging. G. Spatial image plot from sample TRCC6 highlighting distribution of EMT-high tumor cells, EMT-low tumor cells, and SPP1+ macrophages. H. H&E staining of full slide of sample TRCC14. The section highlighted indicates the field of view utilized for spatial imaging.

## SUPPLEMENTARY TABLE LEGENDS

**Table S1.** Clinical characteristics of tRCC sequencing cohort. Related to Figure 1.

**Table S2.** tRCC-specific genes identified from snRNA-seq and bulk RNA-Seq. Related to Figure 2.

**Table S3. Tab 1:** Genes included in each tRCC metaprogram. **Tab 2:** 113 gene fusion activity signature. Related to Figure 3.

**Table S4.** TCRa and TCRb sequences from tRCC cohort profiled in this study. Related to Figure 4 and Figure S4.

